# Enhanced Alcohol-Specific but not General Pavlovian-to-Instrumental Transfer in Alcohol Use Disorder

**DOI:** 10.64898/2026.09.01.747514

**Authors:** Hao Chen, Christian Bäuchl, Louis Thill, Matthew J. Belanger, Michael Marxen, Samanda Krasniqi, Maria Garbusow, Eva Friedel, Maximilian Pilhatsch, Andreas Heinz, Michael N. Smolka

**Affiliations:** Department of Psychiatry and Psychotherapy, Technische Universität Dresden, Dresden, Germany; Department of Sociology, Social Policy, and Criminology, Faculty of Social Sciences, University of Stirling, Stirling, UK; Department of Psychiatry and Psychotherapy, Charité – Universitätsmedizin Berlin, Campus Charité Mitte, Berlin, Germany; Department of Psychology, MSB Medical School Berlin, Berlin, Germany; Department of Psychiatry and Psychotherapy, Klinikum Dresden (SKDF), Dresden, Germany; Department of Psychiatry and Psychotherapy, University of Tübingen and German Center for Mental Health (DZPG), Tübingen, Germany

## Abstract

**Background:** Pavlovian-to-instrumental transfer (PIT) paradigms assess how environmental cues influence instrumental behaviour. Our previous research linked PIT to alcohol use disorder (AUD) and high-risk drinking using monetary rewards during learning. To better capture alcohol-related cue effects, we developed a PIT paradigm delivering trial-by-trial alcohol and juice rewards alongside monetary rewards, to examine alcohol-specific and general PIT effects in parallel.

**Methods:** Seventy-five participants with AUD and ninety-five controls completed the task during functional magnetic resonance imaging. Alcohol-specific PIT was defined as increased alcohol choices during presentation of alcohol-associated versus juice-associated cues. General PIT was defined as increased response vigor during gain-versus loss-associated cues. Behavioural and neural associations with AUD status and AUDIT scores from baseline through one-year follow-up were examined.

**Results:** Participants with AUD showed stronger alcohol-specific PIT effect than controls. Alcohol-specific PIT was positively associated with baseline AUDIT scores and with AUDIT scores across the one-year follow-up. Alcohol cues elicited stronger anterior insula responses, and region-of-interest analysis showed greater left amygdala responses to alcohol versus juice cues in AUD compared with controls. In contrast, general PIT showed a negative association with AUDIT but no association with AUD status. In the general PIT, no AUD-related neural differences were observed.

**Conclusions:** These findings suggest that alcohol-specific PIT captures a clinically meaningful mechanism of cue-driven alcohol seeking that is distinct from generalized motivational transfer, supporting its potential utility as a mechanistic marker in AUD.

## Introduction

Environmental cues significantly shape our daily behaviours. A notification sound on a phone may prompt us to check it immediately, while the smell of freshly brewed coffee may induce a craving in many office workers even though they are not tired. Through Pavlovian learning, neutral cues can acquire motivational value when associated with rewarding experiences or punishments, greatly influencing our ongoing behaviours while minimizing cognitive load. However, reliance on cues can also lead to maladaptive behaviour. For individuals with alcohol use disorder (AUD), alcohol-related cues, such as the sound of opening a beer bottle, can serve as potent triggers and promote alcohol-seeking. This mechanism, where learned cues affect action selection and motivation, can be studied in the laboratory using a Pavlovian-to-Instrumental Transfer (PIT) task (see refs. 1, 2 for reviews).

It is well established that there are two forms of PIT with distinct underlying neural substrates (3, 4). In outcome-specific PIT, the outcome identity signalled by the Pavlovian cue enhances the instrumental action to attain this outcome (e.g. ordering coffee in a café is more likely than ordering alcohol). By contrast, general PIT reflects a broader motivational influence, whereby a positively or negatively valenced cue invigorates or inhibits instrumental actions towards a different rewarding outcome, even when that outcome is not specifically associated with that particular cue (e.g., a lively social atmosphere may encourage drinking).

The theoretical interpretation of outcome-specific PIT remains debated (see ref. 5 for a review). However, increasing evidence suggests that it reflects a goal-directed process (5). From this perspective, individuals actively infer which outcomes are currently likely based on Pavlovian cues and select instrumental actions accordingly (5–9). More broadly, incentive sensitization theory (10–12) proposes that repeated drug use can increase the incentive salience of drug-associated cues, causing them to elicit excessive reward “wanting” even when subjective pleasure (‘liking’) from the reward has diminished. It has been proposed that drug cues may initially evoke outcome-specific transfer effects; however, with repeated drug use, these cues can generalize in their motivational impact, thereby shifting from specific to general transfer effects (13). A key unresolved issue is whether maladaptive cue-driven behaviour in AUD reflects alcohol-specific associative mechanisms or generalized motivational transfer. Previous PIT paradigms have not allowed these processes to be disentangled.

Our research group has established a single-lever PIT task that uses monetary outcomes as reinforcers during both the instrumental and Pavlovian phases, demonstrating links between PIT and alcohol use. The PIT effects observed in this task have been linked to high-risk drinking (14, 15), AUD (16–18), and future relapse (19). However, since participants respond with a single key during the instrumental phase, the task lacked distinct response-outcome mappings, making it unclear whether the observed effects reflect specific or general PIT. Such designs may primarily capture general PIT, as outcome identity does not need to be represented in detail (2). Thus, while these findings suggest that the interactions of Pavlovian and instrumental control are relevant to alcohol use and related disorders, they do not directly address alcohol-specific PIT processes, i.e., the impact of alcohol-related cues on alcohol-seeking behaviour.

To date, three empirical studies have investigated alcohol-specific PIT in association with alcohol-related outcome measures. In all three studies, alcohol-related images or symbols (i.e., pictures of drinks or alcohol points) served as cue reinforcers. Martinovic et al. (20) found no association between the alcohol-specific PIT effect and drinking behaviour in social drinkers. By contrast, Hardy et al. (21) found that alcohol-conditioned stimuli (CSs) increased alcohol choice relative to food choice, and that this effect was positively associated with Alcohol Use Disorders Identification Test (AUDIT) scores in a non-clinical sample of occasionally drinking students. Mahlberg et al. (22) further showed that stronger alcohol-specific PIT was associated with the explicit belief that alcohol was more likely to be received when alcohol cues were present, but not with self-reported craving, supporting the idea that alcohol-specific PIT may partly reflect learned expectations about alcohol availability. Overall, in nonclinical populations, associations between alcohol-specific PIT and alcohol-related measures remain inconsistent. In a related clinical study, van Timmeren et al. (23) assessed both specific and general PIT effects in recently detoxified AUD patients. The task used by this group assessed food-specific and not alcohol-specific PIT, using images of food rather than alcoholic beverages and food pictures as reinforcers. No significant group differences were found between patients with AUD and the control group at either the behavioural or neural level.

The mixed findings across studies may partly reflect a key methodological limitation: previous PIT studies rarely use alcohol-specific PIT tasks and never, so far, actual consumption of alcohol as an immediately available outcome. This may have limited motivational engagement compared to real-life alcohol-seeking behaviours. To improve ecological validity, we developed a novel full-transfer PIT task (24). During the instrumental learning phase, participants received actual gustatory rewards (alcohol and juice). In the Pavlovian conditioning phase, both gustatory outcomes (alcohol and juice) and monetary gains and losses (±€10) were introduced, allowing alcohol-specific PIT effects to be assessed from the gustatory cues and general PIT from the motivational influence of monetary cues on overall response rate. We have shown that this paradigm can elicit both alcohol-specific and general PIT effects in healthy individuals (24). The current study aims to clarify how alcohol-specific and general PIT processes and their neural correlates are associated with AUD and alcohol-related measures.

In the present study, we hypothesized that individuals with AUD would exhibit enhanced alcohol-specific and general PIT at the behavioural level in comparison to individuals without AUD. Smoking is highly prevalent in AUD and may also influence PIT, making it both theoretically relevant and a potential confound. Previous findings are mixed: some studies reported amplified PIT among smokers (e.g., 25), whereas others found no such effect (e.g., 26). Therefore, recruitment in both groups was stratified to balance smoking behaviour across groups to allow examination of smoking-related effects while reducing potential confounding of AUD associations. At the neural level, we expected specific and general PIT to engage regions commonly identified in previous PIT studies, including the amygdala, ventral striatum (VS), putamen, and ventromedial prefrontal cortex (vmPFC) (23, 27–30). Based on our previous findings from monetary PIT paradigms (15, 16, 19), we expected that the ventral striatum and amygdala would exhibit group differences between AUD patients and control participants.

## Materials and Methods

### Participants & Procedure

Participants were recruited in Dresden, Germany, as part of the DFG-funded TRR265 consortium “Losing and Regaining Control over Drug Intake” (31). The study was approved by the Ethics Committee of Technische Universität Dresden (EK 512122018), and all participants provided written informed consent. Recruitment was conducted via flyers and other public advertisements, followed by a telephone screening and in-person assessments. Eligible individuals were 18–60 years old, were fluent in written and spoken German, had no magnetic resonance imaging (MRI) contraindications, and had normal or corrected-to-normal vision. Participants were excluded if they had a lifetime diagnosis of bipolar or a schizophrenia spectrum disorder, substance use disorder other than alcohol, nicotine, or cannabis, a current major depressive episode or suicidal tendency, head injury or severe neurological disease such as dementia, Parkinson’s disease, or multiple sclerosis, pregnancy or breastfeeding, or recent use of medication known to affect the central nervous system.

To balance smoking status, participants were grouped according to a 2 × 3 design ([AUD vs. non-AUD] × [non-smoker, occasional smoker, daily smoker]), with a target of 30 participants per cell (planned N = 180). In total, 190 participants completed the task, of whom 170 remained after exclusions. Individuals assigned to the AUD group were required to meet at least two DSM-5 criteria (32) for AUD based on a structured clinical interview (33), whereas non-AUD participants met fewer than two criteria. Smoking status was assessed using a quantity–frequency questionnaire (items provided in Supplementary Material S-1), defining non-smokers as those who had not smoked in the past 12 months, occasional smokers as those who had smoked in the past 12 months but not daily in the past three months, and daily smokers as those who had smoked seven days per week in the past three months.

Participants first completed a baseline assessment including diagnostic interviews for AUD, tobacco use disorder (TUD), and other selected psychiatric disorders, neuropsychological tests, and questionnaires including a quantity-frequency questionnaire for alcohol use (34), a quantity-frequency questionnaire for smoking, the AUDIT (35), the Craving Automated Scale (CAS) (36) and the Fagerström Test for Nicotine Dependence (FTND) (37). Those meeting the eligibility criteria were invited to a first MRI session within four weeks, during which they performed a single-lever PIT task; data from this task are not included in the present analyses. Approximately one week later, they returned for the full-transfer PIT task.

Following the baseline assessment, participants completed three online follow-up assessments over 12 months, focusing primarily on drinking behavior. The measure of interest at follow-up was the AUDIT, which assessed alcohol-related behavior since the previous assessment.

### Full-transfer PIT task

Before the task, participants sampled six alcoholic beverages and six fruit juices and rated their pleasantness on a 7-point scale. For each participant, one alcoholic drink and one juice were selected that were highly rated in pleasantness and closely matched in pleasantness to minimize value differences between the two rewards. The full beverage list and detailed selection procedure are described in Belanger et al. (24).

The task consisted of four phases (Figure 1): instrumental learning, Pavlovian conditioning, a PIT transfer phase, and query trials to confirm Pavlovian learning. In the instrumental phase, participants learned the association between two response keys and the delivery of the selected alcoholic beverage and juice, with key-drink assignments randomized across participants. On each trial, upon seeing an exclamation mark, participants were instructed to choose one key and press it repeatedly during a 2-second response window. When participants made at least five presses (the response criterion was unknown to participant), the trial was reinforced with a 50% probability through delivery of 1 mL of the corresponding drink intraorally via a syringe pump. Of the reinforced trials, 20% delivered the alternative drink to introduce outcome uncertainty. The instrumental phase continued until each drink had been delivered 12 times, ensuring balanced learning of the action-outcome associations for both alcohol and juice. On average, participants completed the instrumental learning phase in 57 ± 17 trials. Learning of the key–drink associations was confirmed by subsequent action-outcome knowledge check (Figure 1).

**Figure 1:**
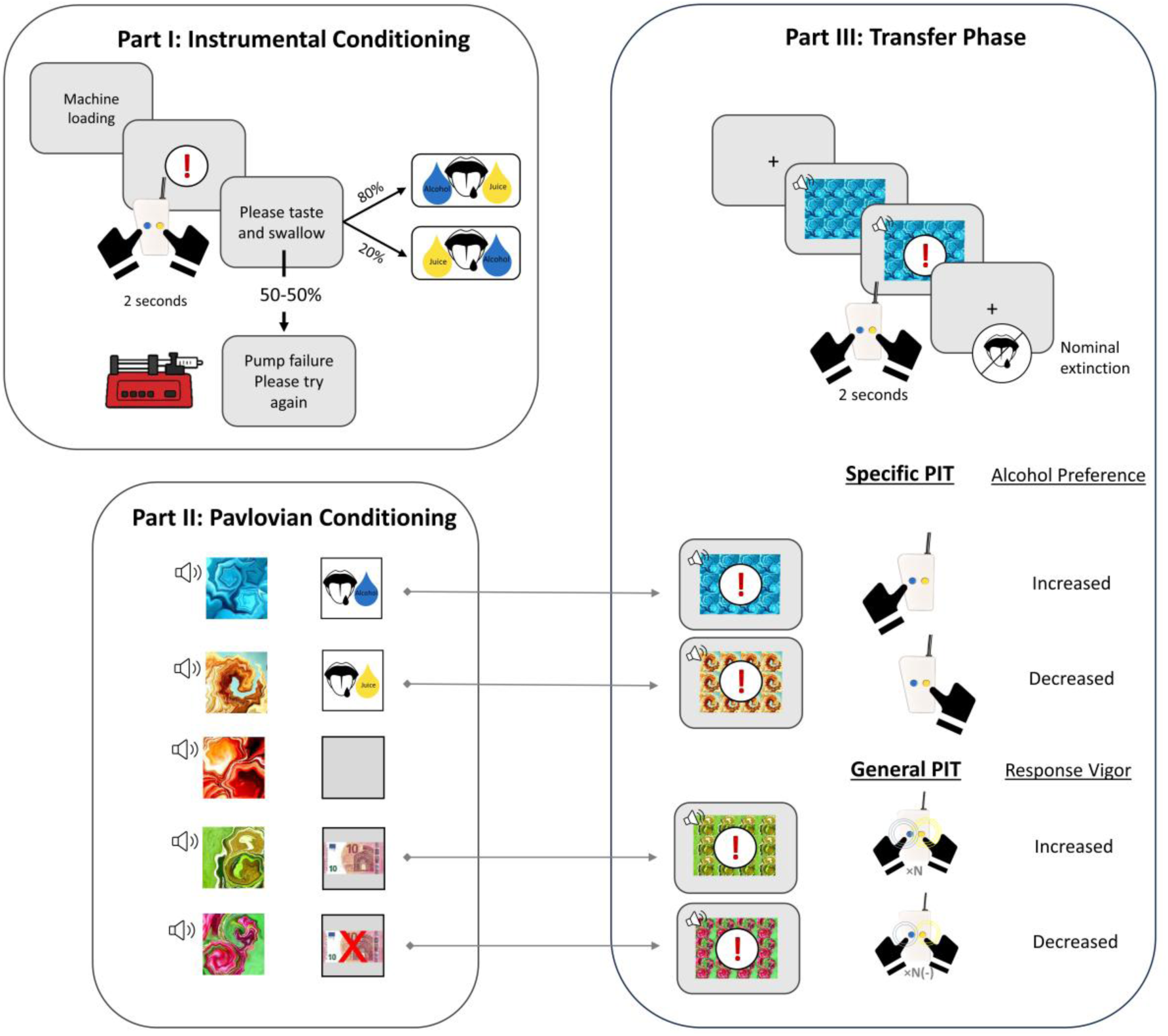
Full Transfer PIT Paradigm. **Figure 1: Schematic of the full transfer task.** The task comprised the following phases: **(I) Instrumental learning:** Participants were instructed to repeatedly press one of two response keys within a 2,000-ms response window to receive an intraoral delivery of alcoholic drink or juice. A minimum of five button presses was required for a potential drink reward. To introduce uncertainty and avoid fully deterministic learning, 50% of the trials were rewarded, while nonrewarded trials were presented as pump failures. On rewarded trials, the selected response led to its associated drink in 80% of cases and to the alternative drink in 20% of cases. The phase ended after participants had received 12 alcohol and 12 juice rewards. Participants completed action-outcome knowledge check assessing the learned key–drink associations. Learning was high, with 97.2% accuracy for the alcohol-associated key and 96.9% for the juice-associated key. **(II) Pavlovian conditioning:** five compound cues (fractals paired with tones) were associated with alcohol, juice, monetary gain, monetary loss, or no outcome. After a 500-ms fixation, cues were presented for 2,500 ms. Gustatory outcomes were delivered during cue presentation, followed by 1,000 ms for tasting and 2,000 ms for swallowing; monetary outcomes were displayed for 1,000 ms after cue presentation. **(III) The transfer phase (fMRI; under nominal extinction):** Participants performed the instrumental response with a particular Pavlovian cue tiled in the background. After a 500-ms fixation, the Pavlovian cue appeared for 600 ms before the 2,000-ms response window and remained visible throughout responding. Alcohol and juice cues were used to assess specific PIT, reflecting whether alcohol-related cues increased selection of the alcohol-associated response relative to juice cues. Monetary gain (€+10) and loss (€-10) cues were used to assess general PIT, reflecting cue-related changes in overall response vigor irrespective of the specific drink outcome.

During the Pavlovian conditioning phase, five compound cues (fractals paired with musical tones) were randomly assigned across participants and consistently associated with one of the following five outcomes: alcohol, juice (via syringe pump), a monetary gain of €10, a monetary loss of €10, or no outcome. Each cue-outcome pairing was presented for 16 trials, resulting in a total of 80 trials.

The PIT transfer phase took place under nominal extinction inside the MRI scanner. During each trial, one of the Pavlovian cues was tiled in the background (see Figure 1) while the musical tone was played through headphones. Participants were instructed to choose freely which key and how many times to press it upon seeing the exclamation mark. No drinks were delivered and no monetary outcomes were presented during the scanner trials. Instead, participants were told that drinks or money would be accumulated and provided after the task. Alcohol– or juice-associated cue were expected to bias responding toward the instrumental action associated with the same outcome (specific PIT). Cues associated with monetary gain or loss, which were unrelated to either instrumental action, were used to assess general transfer effects, i.e., cue-related changes in overall response vigor, indexed by the number of button presses. Each Pavlovian cue was presented 48 times, resulting in 240 trials for the PIT phase.

After the transfer phase, participants completed query trials in which they indicated which outcome was associated with each cue, providing an explicit measure of Pavlovian learning. A short debriefing questionnaire assessed task comprehension. The results from the query trials are detailed in the Supplementary Material S-2.

### MRI data acquisition

MRI data for 48 participants were acquired on a 3-Tesla MAGNETOM Trio Tim and 122 participants on the PrismaFit upgrade of the same scanner (Siemens Healthineers AG, Forchheim, Germany) at the Neuroimaging Center of Technische Universität Dresden using the manufacturer’s 32-channel head coils. Structural 3D T1-weighted magnetization-prepared rapid gradient-echo (MPRAGE) images were obtained (repetition time (TR): 2000 ms, echo time (TE): 2.01 ms, field of view (FOV): 256 mm, flip angle (FA): 8°, time of inversion (TI): 880 ms, voxel dimensions: 1 × 1 × 1 mm). Prior to echo planar imaging (EPI) acquisition, a gradient-echo sequence was recorded to obtain a field map, which was used to correct for geometric distortions caused by static field inhomogeneities (TR: 698 ms, TE 1: 5.19 ms, TE 2: 7.65 ms, FOV: 210 mm, number of slices: 64, voxel dimensions: 2.4 × 2.4 × 2.4 mm). Functional data were acquired using a T2*-weighted EPI sequence with a multiband acceleration factor of 6 (TR: 869 ms, TE: 38 ms, FA: 58°, in-plane FOV: 210 mm, number of slices: 60, voxel dimensions: 2.4 × 2.4 × 2.4 mm). For EPI acquisition, slices were tilted by –25° relative to the anterior commissure-posterior commissure (AC-PC) line.

### Behavioural Analyses

To capture the alcohol-specific PIT effect, we first calculated participants’ preference for the alcohol button during presentation of the alcohol CS and during presentation of the juice CS. Participants who failed to respond on more than 50% of transfer trials were excluded. For the remaining participants, trials with no response or responses for both reward options were excluded from the analysis. The preference for alcohol (*p_alcohol_*) was calculated as the proportion of alcohol choices: *n_alcohol_ / (n_alcohol_ + n_juice_)* (100% = participant always pressed the alcohol button; 0% = participant always pressed the juice button during a particular CS presentation). We then calculated the specific PIT score as the increase in *p_alcohol_* during the presentation of the alcohol CS compared with the juice CS ([*p_alcohol_* | alcohol CS] – [*p_alcohol_* | juice CS]). This difference reflects the extent to which the alcohol and juice CSs bias participants’ preference for alcohol choices: 100% = always alcohol when the alcohol CS is shown and never alcohol when the juice CS is shown; 0% = alcohol preference is the same during presentation of both CSs. Note that negative numbers are unexpected but not impossible. Given the non-normal distribution of the PIT score, a one-sample Wilcoxon signed-rank test was first applied to determine whether there was a significant specific PIT effect at the sample level.

Following recommendations to use robust tests as the default analytic approach (38) and given the non-normal residual distribution in the corresponding ordinary least-squares models, we used robust regression as the primary analytic approach. This approach reduces sensitivity to distributional deviations and limits the influence of extreme values while retaining all observations. Robust regression was implemented using the *rlm* function from the R package *MASS* (39). For robust regression models, we report 95% bias-corrected and accelerated (BCa) bootstrap confidence intervals (CIs) based on 5,000 resamples, the bootstrap standard error (SE) and bootstrap p values. In the initial full model, the specific PIT score was entered as the dependent variable. AUD group, smoker group, and their interaction were included as predictors, with age and sex added as covariates. Smoking group was treated as an ordered linear predictor (non-smoker < occasional smoker < daily smoker) and mean-centred before computing the AUD × smoking interaction; age was also mean-centred. The model was specified as:

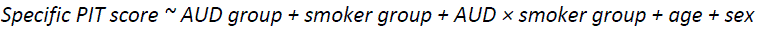

Because smoking group, age, and sex were not associated with the specific PIT score (reported in more detail in the Results section), these variables were not included in the subsequent dimensional analyses. Exploratory dimensional analyses across all participants were conducted using separate robust regression models to examine associations between the specific PIT score and drinking-related measures, including AUDIT total score and number of DSM-5 AUD criteria as measures of AUD severity; the AUDIT consumption subscore (AUDIT-C), alcohol consumption in grams per drinking occasion, and drinking frequency per week as measures of alcohol consumption; and the CAS craving score as a measure of craving.

General PIT effects were captured by analyzing participants’ response vigor, i.e., the number of button presses regardless of choice (alcohol or juice), during the +10€ CS relative to the –10€ CS. In this analysis, trials without a response were not excluded but coded as zero button presses, reflecting low motivation. The general PIT score was calculated as the difference in response vigor (i.e., number of button presses) between the +10€ CS and –10€ CS conditions. Parallel to the statistical test for specific PIT effect, we first performed a parallel robust regression using the same independent variables as for specific PIT:

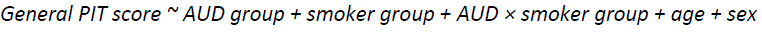

The same exploratory dimensional robust regression analyses with the six drinking-related measures were then repeated for the general PIT score.

To determine whether associations between PIT and alcohol-related problems extended beyond the concurrent baseline assessment, we conducted exploratory analyses examining alcohol-specific and general PIT in association with AUDIT scores across the one-year follow-up period. For these analyses, we fitted robust linear mixed-effects models of AUDIT measures from all available assessment points with the *rlmer* function from the R package *robustlmm* (40). AUDIT total score and AUDIT-C were examined in separate models as dependent variables. Each model included PIT score, time point, and their interaction as fixed effects, with participant-specific random intercepts to account for repeated measurements. Separate models were fitted for specific and general PIT scores. The PIT main effect tested whether PIT was associated with overall drinking severity, whereas the PIT × time interaction tested whether this association varied across assessment points. 95% participant-level bootstrap CIs based on 1,000 resamples, as well as the bootstrap p values were computed for the fixed-effect estimates.

### Functional MRI preprocessing and statistical analyses

#### Preprocessing

Basic preprocessing of imaging data was conducted with SPM12 (Statistical Parametric Mapping; www.fil.ion.ucl.ac.uk/spm). Functional data were slice-time corrected via SPM’s Fourier phase shifting interpolation, using the middle slice as the reference. Functional images were spatially realigned to the first frame and then to the mean image of the experimental run to correct for head motion. Geometric distortions in EPI images were corrected by calculating voxel displacement maps (VDMs) from individual field maps. The individual structural T1-weighted image was coregistered to the mean EPI image and segmented into gray and white matter. Both functional and structural images were normalized to the standard space of the Montreal Neurological Institute (ICBM 152 MNI template) based on the T1-derived transformation obtained from the segmentation and resampled to their original voxel resolution during acquisition at 2.4 mm³ and 1 mm³, respectively. Finally, functional images were spatially smoothed with an isotropic 8-mm full-width-at-half-maximum (FWHM) Gaussian kernel.

#### First-level analysis

For subject-level BOLD analyses of the PIT phase, task-related fMRI time series were modeled in a general linear model (GLM) with regressors convolved with the canonical hemodynamic response function. Regressors of interest comprised (i) alcohol and juice CS onset regressors, each parametrically modulated by the choice (coded as +1 for alcohol choice and −1 for juice choice), and (ii) −€10, neutral, and +€10 CS onset regressors, each parametrically modulated by the total number of button presses during the corresponding CS presentation (i.e., alcohol and juice presses combined). To mitigate the effects of in-scanner head motion, the following regressors were additionally included: a Volterra expansion of the six motion parameters (41) and separate censoring regressors for each scan with FD > 0.9 (42, 43) as covariates of no interest in the design matrix. The Volterra expansion comprised linear and quadratic effects of the estimated motion parameters (from spatial realignment) and linear and quadratic effects of the first derivatives of said parameters. Together, this resulted in 24 covariates which are able to capture higher-order effects of motion, like spin-history effects (44). For motion censoring, we first calculated the framewise displacement (FD) of every volume of each individual’s time series according to Power et al. (42). Then, we set the criterion for excessive head motion to FD > 0.9 (43) and created scan nulling regressors that flagged affected scans, thereby removing their influence on the parameter estimates of interest. Finally, functional data were high-pass filtered with a cutoff of 128 s and corrected for temporal autocorrelation using the FAST model.

At the first level, we defined a specific cue contrast comparing the alcohol CS and juice CS onset regressors (Alcohol CS > Juice CS) to quantify differential neural responses to alcohol versus juice cues during the PIT phase (i.e., a cue-reactivity component relevant to the specific PIT). In addition, we tested a choice-modulated specific PIT contrast based on the choice by comparing neural responses associated with choosing alcohol over juice during alcohol CS presentations with those associated with choosing alcohol over juice during juice CS presentations, i.e., the difference of the choice parametric regressors between the two CS types. This contrast tested whether alcohol cues preferentially bias alcohol-versus-juice choice activations at the neural level, consistent with a within-subject specific PIT effect described by van Timmeren et al. (23).

Similarly, a general PIT cue contrast compared +€10 and −€10 CS onsets. A vigor-modulated general PIT contrast compared the corresponding button-press parametric regressor coefficients for +€10 versus −€10 trials, testing whether BOLD responses scaled more strongly with response vigor during the presentation of appetitive (+€10) than aversive (−€10) cues.

#### Second-level whole-brain analysis

These first-level contrast images were then entered into second-level analyses to estimate sample-level effects for both specific and general PIT. The main whole-brain analyses focused on first-level cue contrasts: the alcohol CS vs. juice CS contrast for specific PIT and the +€10 CS vs. −€10 CS contrast for general PIT. In addition, two separate between-subject brain-behavior regression analyses tested whether individual behavioral PIT effects (specific PIT and general PIT scores), entered as covariates on the second level, were associated with the corresponding cue-related neural responses. This approach tested whether inter-individual variability in behavioral PIT is associated with the magnitude of cue-related neural responses.

Furthermore, to examine differences between AUD and non-AUD participants, we conducted two-sample *t*-tests on the specific PIT contrast and the general PIT contrast. Whole-brain analyses of *within-subject* PIT effects based on choice-/vigor-related parametric regressors were conducted as exploratory analyses and are reported in the Supplementary Material S-3. Whole-brain statistical maps were thresholded at p < .001 (uncorrected) with a minimum cluster extent of k > 50 voxels.

#### Region of interest (ROI) analysis

Following the whole-brain analysis, we conducted region of interest (ROI) analyses to test whether cue-related neural responses differed between groups and whether they were associated with behavioral PIT parameters. Based on the previous PIT literature (23, 27–30), we selected the amygdala, VS, putamen, as well as vmPFC as ROIs.

The bilateral amygdala and putamen ROIs were defined anatomically using the Automated Anatomical Labeling (AAL) atlas (45) as implemented in the WFU PickAtlas toolbox (46, 47). The vmPFC ROI was derived from the Neurosynth (https://www.neurosynth.org/) meta-analytic map by querying the term “vmPFC”; voxels assigned to the anterior cingulate cortex were then excluded. The ventral striatum (VS) mask was obtained from the BrainMap database (48) by searching for the term “accumbens”. The meta-analytic ROIs were then spatially smoothed and binarized; the VS mask was removed from the vmPFC mask to prevent overlap. All ROIs were resliced to 2.4 × 2.4 × 2.4 mm to match the functional data resolution.

For each ROI, we extracted mean contrast estimates for the specific and general PIT cue contrasts. We first compared ROI responses between the AUD and non-AUD groups and then examined dimensional associations with AUDIT total score and AUDIT-C. Here, we used ordinary linear regression, as model assumptions were met. Following this, to test whether neural cue responses were related to individual differences in PIT, we examined associations between ROI responses and behavioral indices of specific and general PIT effects using robust regression with confidence intervals, consistent with the behavioral analyses, because models involving the non-normally distributed PIT behavioral parameters showed less optimal model assumption diagnostics.

Since we did not find any behavioral associations between smoking status and PIT, and given the limited sample size for additional subgroup imaging analyses, we did not extend the imaging analyses to smoking status.

## Results

### Sample characteristics

A total of 190 participants completed the full-transfer PIT task. One participant was excluded due to an experimenter error during pump operation. Eighteen participants were excluded because they did not respond in more than 50% of trials, and one additional participant was excluded for falling asleep during the task. Among the remaining 170 participants, 2,480 no-response trials (6.0% of 41,040) and 180 trials with simultaneous left– and right-button presses (0.4%) were further removed.

The final sample comprised 75 participants with AUD and 95 non-AUD controls (see Table 1). Most participants in the AUD group had mild-to-moderate AUD (79%; 2-5 DSM-5 criteria). The AUD group was significantly older than the non-AUD group (mean age: 34.5 ± 12.7 vs. 29.8 ± 8.0 years, p = .006). The groups did not differ in sex distribution, monthly income, or educational status. As expected, the AUD group showed higher AUD severity and higher AUDIT scores. The AUD group also reported higher alcohol consumption (81.2 ± 42.4 vs. 31.0 ± 20.1 g/drinking day, p < .001) and higher frequency of drinking (4.3 ± 1.7 vs. 1.0 ± 1.1 drinking days/week, p < .001) during the last three months. Due to the sampling strategy, there were no differences in the distribution of smokers, TUD criteria, or FTND scores between the two groups. Descriptive statistics and statistical tests results are detailed in Table 1. For the follow-up drinking assessment, 107 participants completed at least one follow-up AUDIT questionnaire (68 with AUD and 39 non-AUD participants).

**Table 1:**
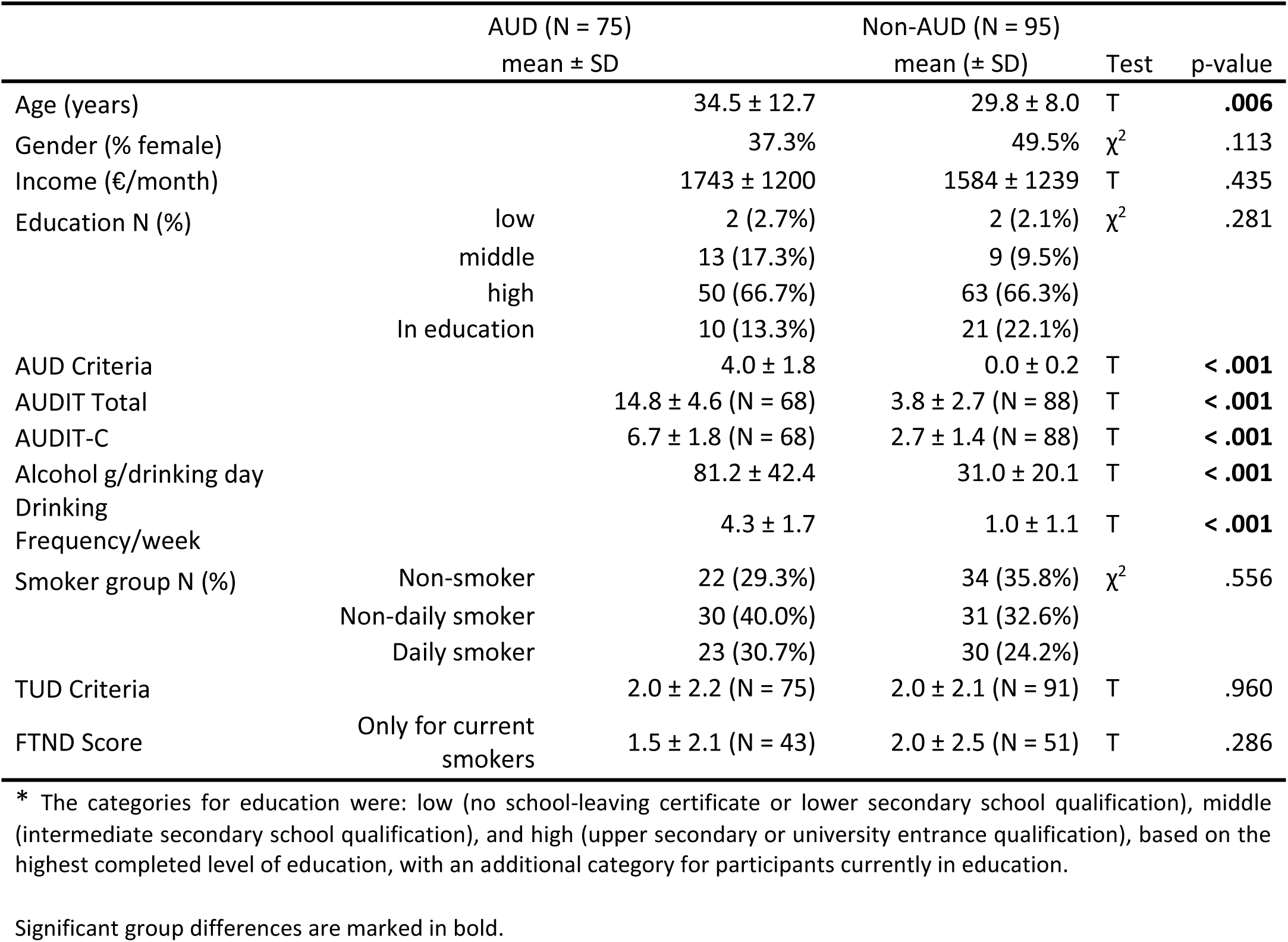
Sample Characteristics.

### Behavioral Results

#### Specific PIT: cross-sectional associations

With respect to specific PIT, positive values indicate that the presentation of alcohol CS increases preference for alcohol, whereas negative scores indicate a decrease in preference. We calculated the specific PIT score as the difference in the proportion of alcohol choices between the alcohol CS and juice CS conditions (% alcohol choice | alcohol CS – % alcohol choice | juice CS). On the group level, the specific PIT score was significantly greater than zero (Wilcoxon signed-rank test: p < .001; r = 0.702), indicating an overall increase in alcohol choices during the presentation of alcohol compared with juice cues (increase from 9.9% to 56.1%; Figure 2A).

**Figure 2:**
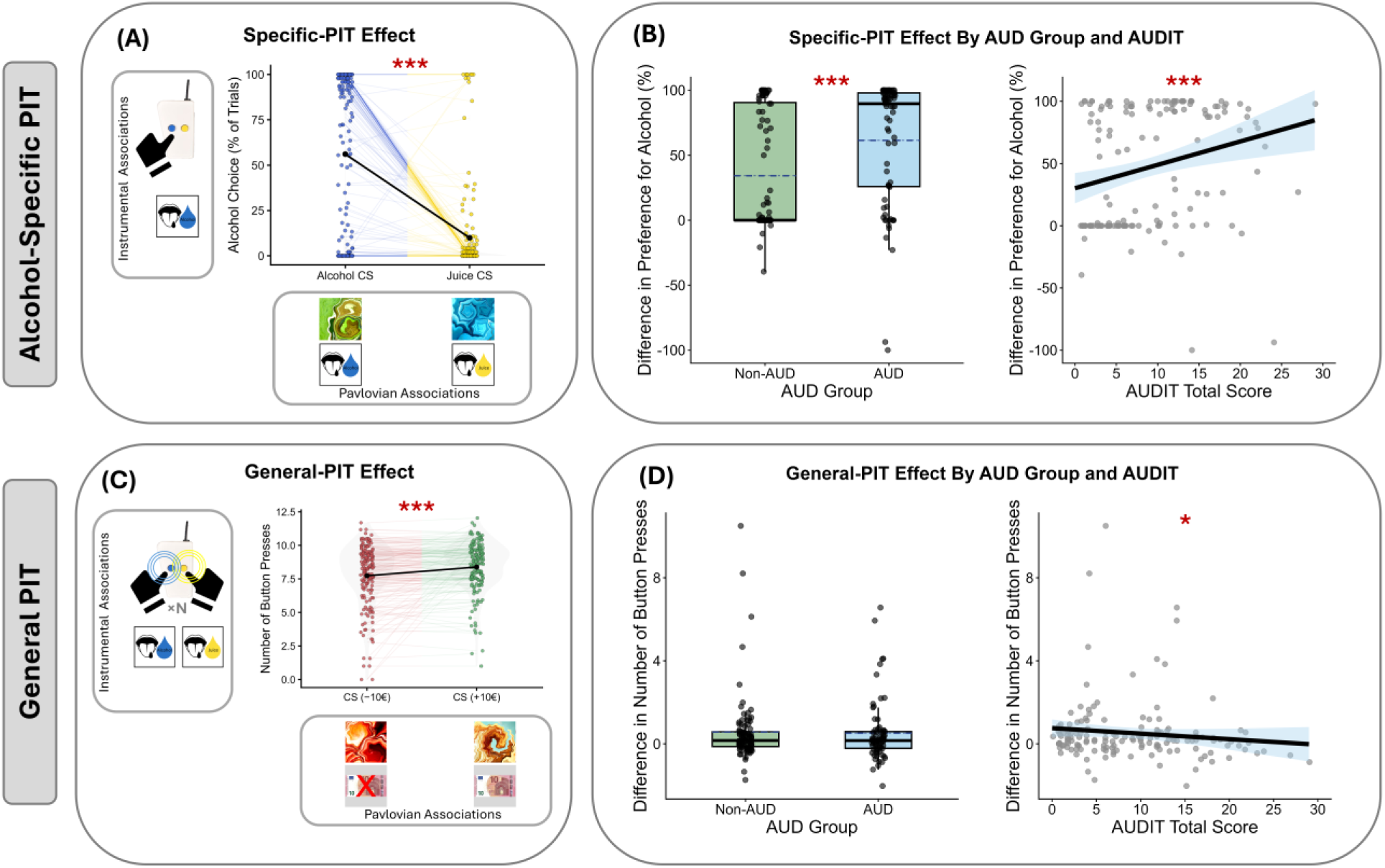
Behavioural PIT effects. **Figure 2: Behavioural PIT effects. (A)** At the sample level, a pronounced alcohol-specific PIT effect was observed: participants made more alcohol choices when presented with an alcohol cue (56.1%) than when presented with a juice cue (9.9%). **(B) Left:** The specific PIT effect was significantly higher in the AUD group (p < .001). The distribution of the specific PIT effect (i.e., the difference in preference for alcohol) is shown for the Non-AUD and AUD groups, with individual data points. The dashed blue line indicates the mean, and the boxes show the interquartile range with the median line. **Right:** The specific PIT effect demonstrated a positive association with the AUDIT total score (p < .001); the light blue area represents the 95% confidence interval. **(C)** A significant general PIT effect was observed at the sample level (p < .001), with 0.6 more button presses in the +€10 condition than in the –€10 condition. **(D) Left:** There was no group difference in general PIT effect (denoted as the difference in number of button presses in the +€10 Pavlovian cue condition compare with –€10 condition; p = .315). The dashed blue line indicates the mean, and the boxes show the interquartile range with the median line. **Right:** The association between the general PIT effect and AUDIT total score was negative (p = .012); the light blue area represents the 95% confidence interval. Significance levels are indicated by p < .05 (*) and p < .001 (***).

In the full robust regression model, we examined whether specific PIT scores differed by AUD group while accounting for smoker group, the AUD × smoker interaction, age, and sex. Smoking group and age were mean-centered, such that the AUD group coefficient reflected the AUD versus non-AUD difference at the average smoking level and average age. Specific PIT scores were higher in the AUD group than in the non-AUD group, β = 30.88, SE = 9.90, p < .001, 95% bootstrap CI [15.47, 54.65] (displayed in Figure 2B). Smoker group was not associated with specific PIT scores, β = 7.43, SE = 6.03, p = .193, CI [−4.54, 19.03]. The AUD × smoker group interaction was not supported either, β = −9.36, SE = 9.33, p = .315, CI [−26.69, 9.73]. Age and sex were not associated with specific PIT scores, with bootstrap CIs including zero (full results in Table 2).

**Table 2:**
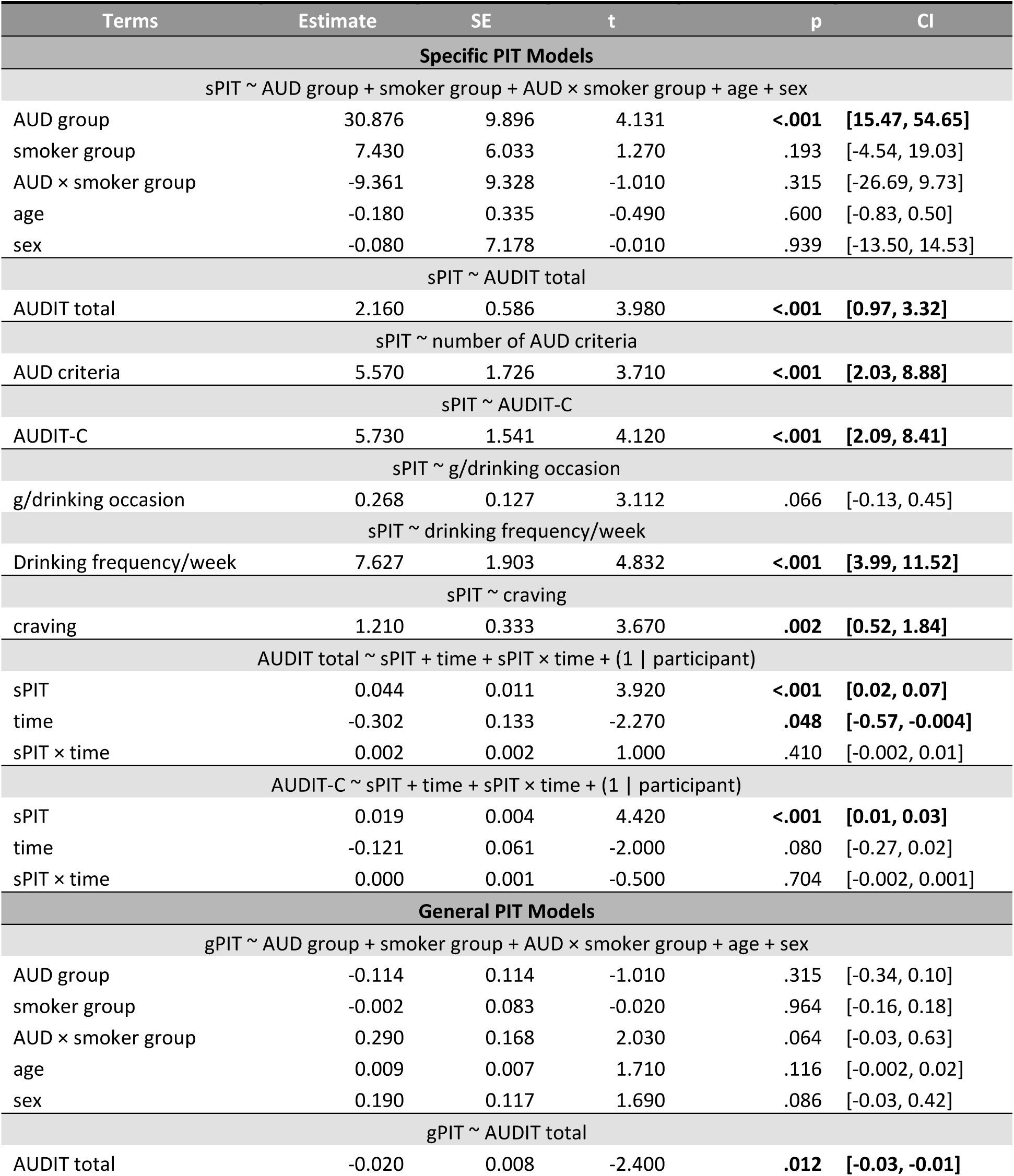
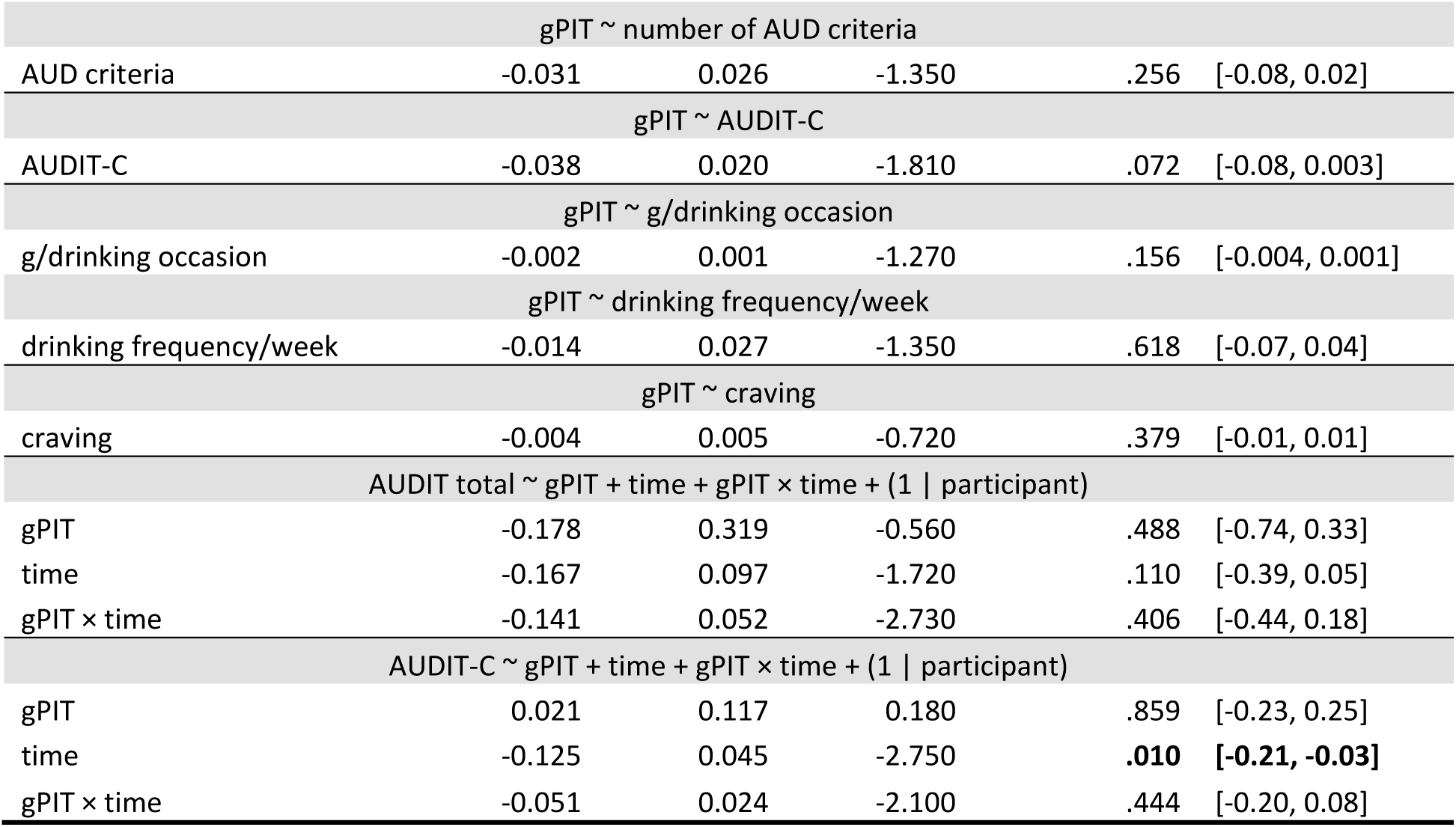
Full results from regression models.

In the dimensional robust regression analyses (which did not include age, sex and smoking status), drinking-related measures showed consistent positive associations with specific PIT scores. Regarding AUD severity, higher AUDIT total scores were associated with higher specific PIT scores, β = 2.16, SE = 0.59, p < .001, CI [0.97, 3.32] (displayed in Figure 2B); similar positive associations were found for the number of AUD criteria (β = 5.57, SE = 1.73, p < .001, CI [2.03, 8.88]). Regarding consumption, positive associations were observed for AUDIT-C, β = 5.73, SE = 1.54, p < .001, CI [2.09, 8.41], and for drinking frequency (β = 7.63, SE = 1.90, p < .001, CI [3.99, 11.52]), but marginal for drinking quantity (gram/drinking occasion, β = 0.27, SE = 0.13, p = .066, CI [-0.13, 0.45]). Moreover, craving scores were also positively associated with specific PIT scores (β = 1.21, SE = 0.33, p = .002, CI [0.52, 1.84]).

#### Specific PIT: association with follow-up drinking assessments

To test whether alcohol-specific PIT has a trait component rather than reflecting merely current drinking behaviour, we next examined its association with longitudinal AUD measures. In more detail, we examined whether specific PIT effect was associated with AUD severity (AUDIT) and consumption (AUDIT-C) across baseline and the one-year follow-up assessments with robust mixed-effects models. We found that higher specific PIT scores were associated with higher AUDIT total scores across all time points, β = 0.044, SE = 0.011, p < .001, 95% participant-level bootstrap CI [0.02, 0.07], and higher AUDIT-C scores, β = 0.019, SE = 0.004, p < .001, CI [0.01, 0.03]. The specific PIT × time interactions were not significant for either AUDIT total, β = 0.002, SE = 0.002, p = .410, CI [-0.002, 0.01], or AUDIT-C, β = −0.001, SE = 0.001, p = .704, CI [-0.002, 0.001], indicating that these associations were not limited to baseline but remained relatively stable over time.

#### General PIT: cross-sectional associations

General PIT was quantified as the difference in response vigor (number of button presses) between the +€10 and –€10 conditions. Overall, participants pressed the button more often in the +€10 condition (p < .001, r = 0.357) than in the –€10 condition, with an average difference of 0.6 button presses, indicating increased response vigor (Figure 2C).

In the categorical robust regression model for general PIT, AUD group did not affect general PIT scores, β = −0.11, SE = 0.11, p = .315, 95% CI [−0.34, 0.10] (Figure 2D). Smoker group was also not associated with general PIT scores, β = −0.002, SE = 0.08, p = .964, CI [−0.16, 0.18]. The AUD × smoker group interaction showed a marginal positive effect, β = 0.29, SE = 0.17, p = .064, and the bootstrap CI including zero, CI [−0.03, 0.63]. Age and sex were not associated with general PIT scores (Table 2).

Regarding the dimensional drinking measures, general PIT scores were negatively associated with AUDIT total score, β = −0.020, SE = 0.008, p = .012, CI [−0.03, −0.01] (Figure 2D) but not with the number of AUD criteria (β = −0.031, SE = 0.026, p = .256, CI [−0.08, 0.02]). AUDIT-C showed a trend-level negative association, β = −0.038, SE = 0.020, p = .072, CI [−0.08, 0.003]. No associations were found for gram/drinking occasion, β = −0.002, SE = 0.001, p = .156, CI [−0.004, 0.001], drinking frequency, β = −0.014, SE = 0.027, p = .618, CI [−0.07, 0.04], or craving, β = −0.004, SE = 0.005, p = .379, CI [−0.01, 0.01].

#### General PIT: association with follow-up drinking assessments

When examining the associations between general PIT and drinking measures across the follow-up assessments, we found no evidence for an overall association between general PIT and AUDIT total or AUDIT-C scores across the assessment period. The general PIT × time interaction was not significant for either the AUDIT total or AUDIT-C (results detailed in Table 2).

### fMRI Results (1177 words)

#### Whole-brain results: specific-PIT effect

Eight participants were excluded from the fMRI analysis because of acquisition or preprocessing issues: three because the task was not completed in the scanner, one because task onset was not synchronized with scan triggering, two due to missing field maps, one due to a technical issue affecting the field map data, and one because the BOLD data were improperly recorded due to a technical error. The final imaging sample comprised 162 participants.

We first examined whole-brain responses to alcohol-specific Pavlovian cues and their association with individual differences in the behavioral alcohol-specific PIT effect (between-subject brain-behaviour analysis). For the specific PIT cue-related contrast (alcohol CS − juice CS), at a whole-brain threshold of p_uncorr_. <.001 and k > 50, greater fMRI responses were observed in a distributed network (Figure 3A). These included the left insula (k = 554; peak MNI coordinate: – 37/15/2; t = 4.47) and orbitofrontal cortex (k = 552; peak MNI coordinate: 42/22/-3; t = 4.79), lateral prefrontal cortex (bilateral middle frontal gyrus; peaks: 40/20/45 and −37/15/43), as well as temporo-parietal association regions including the angular gyrus, precuneus, posterior cingulate, and posterior temporal cortex (detailed in Table S1). In a single cluster in the left amygdala (k = 80; peak MNI coordinate: –23/-9/-15; t = 4.07) fMRI responses were lower during alcohol CS than juice CS presentation (Figure 3B).

**Figure 3:**
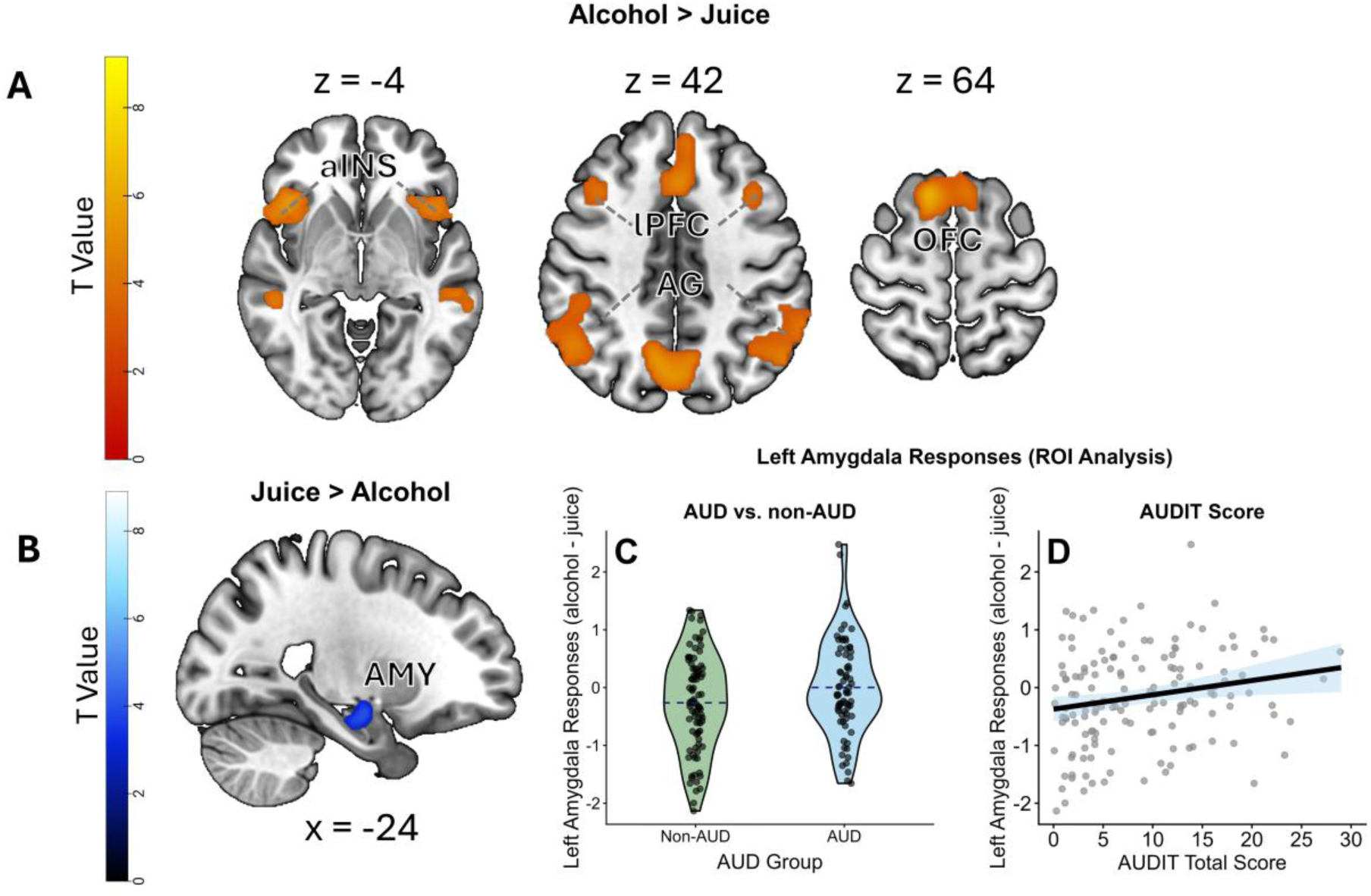
Neural associations for Specific-PIT Effect. **Figure 3: Neural associations for specific PIT effect. (A)** Neural responses for the alcohol > juice cue contrast, including anterior insula (aINS), lateral prefrontal cortex (lPFC), angular gyrus (AG) and orbitofrontal cortex (OFC). (B) Neural responses during the juice > alcohol cue contrast in the left amygdala **(C)** AUD group showed more differences in the left amygdala neural responses during the alcohol cue compared to the juice cue presentation. The dashed blue line indicates the mean. **(D)** The differences in the left amygdala neural responses were positively associated with the AUDIT total score across all participants. The light blue area represents the 95% confidence interval.

For the between-subject brain-behavior analysis of the alcohol-specific effect, higher PIT scores were associated with greater neural responses in the left postcentral gyrus (k = 96; peak MNI coordinate: –61/-16/31; t = 4.19) and anterior cingulate cortex (ACC; k = 71; peak MNI coordinate: 8/39/7; t = 4.05). In contrast, only one cluster in the lateral occipital cortex showed a negative association with the alcohol-specific PIT score (Table S1).

#### Whole-brain results: general-PIT effect

We next examined whole-brain responses to monetary Pavlovian cues and their association with the behavioural general PIT effect. The +€10 > −€10 cue contrast engaged a large reward-related network, dominated by a large cluster peaking in the left putamen (k = 17181; peak: −30/−6/−10, t = 5.99) extending into widespread cortical regions including posterior cingulate/precuneus, lateral occipital cortex, posterior temporal cortex, precentral/SMA, thalamus, and angular gyrus, as well as the amygdala and hippocampus (Figure 4A). Additional clusters were found in the right medial frontal cortex (k = 4063; peak: 6/46/−10, t = 5.36), left orbitofrontal cortex (k = 151; peak: −47/27/−17, t = 4.60), left precentral gyrus, right temporal pole, and bilateral cerebellum (Table S1). The reverse contrast (-€10 > +€10) yielded no significant clusters.

**Figure 4:**
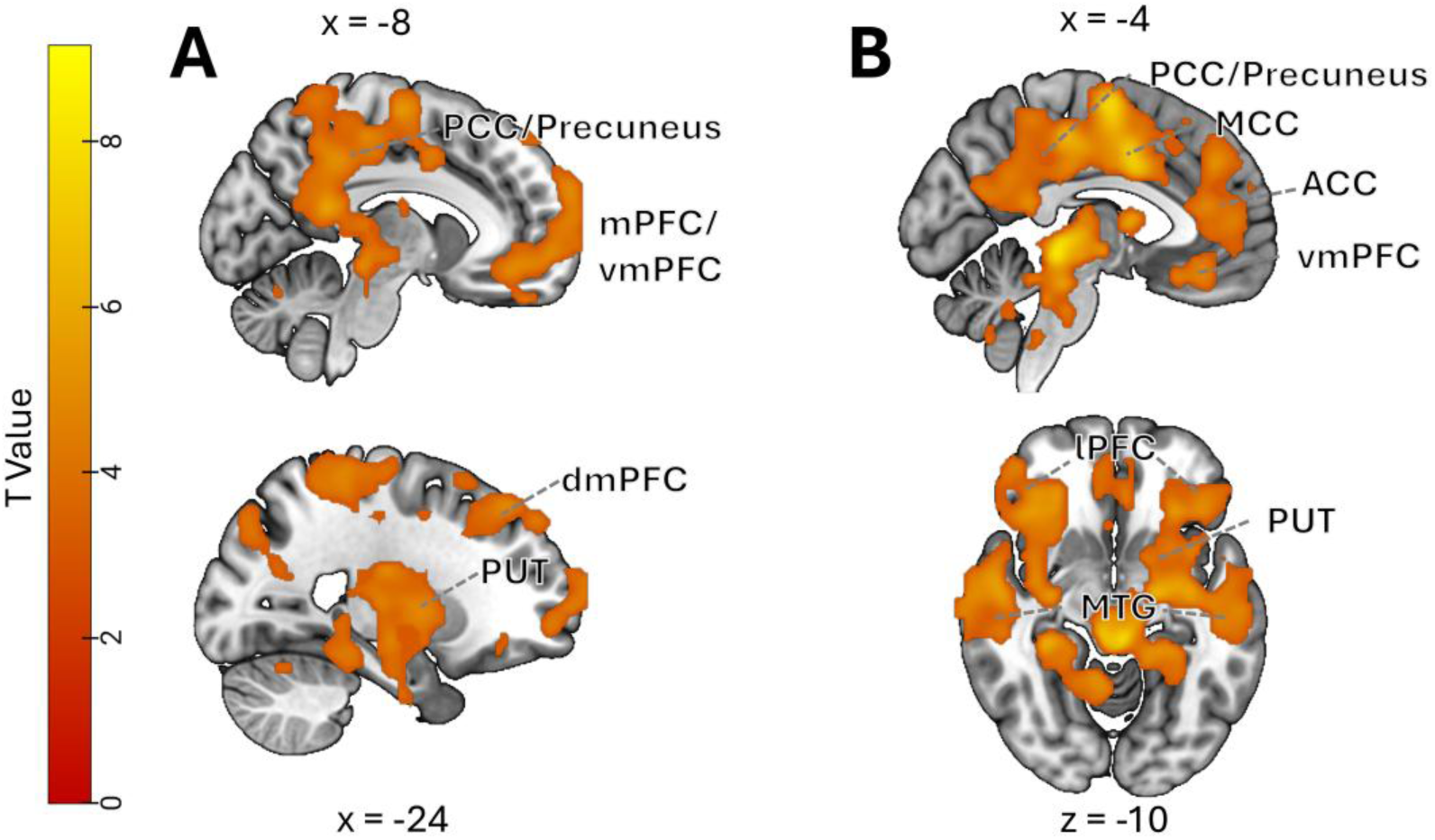
General PIT neural responses. **Figure 4: Neural responses and between-subject associations for general PIT effect.** (A) Neural responses when for the +€10 > −€10 cue contrast, including posterior cingulate cortex (PCC), precuneus, ventromedial/medial prefrontal cortex (mPFC/vmPFC), dorsomedial prefrontal cortex (dmPFC) and putamen (PUT). (B) Neural responses for the +€10 > −€10 cue contrast associated with individual differences in behavioral general PIT, including PCC, precuneus, mid-cingulate cortex (MCC), anterior cingulate cortex (ACC), vmPFC, lateral prefrontal cortex (lPFC), middle temporal gyrus (MTG) and PUT.

The between-subject brain-behaviour general PIT analysis yielded a large cluster with a peak in the medial cingulate/premotor region (k = 43,838; peak: –8/3/38, t = 9.27), with prominent involvement of the basal ganglia (mainly putamen, but also pallidum and caudate), and extended into cingulate cortex (anterior to posterior), insula, motor/premotor cortex, frontal cortex (left superior/middle frontal and right orbitofrontal/frontopolar regions), and brainstem (Figure 4B; Table S1). Negative associations with the general PIT score were observed in right cerebellar crus and right occipital pole.

For the group differences, we first conducted two-sample *t*-tests on the whole-brain alcohol CS > juice CS contrast (specific PIT) and +€10 > –€10 CS contrast (general PIT). At the prespecified whole-brain threshold, no significant differences between the AUD and non-AUD groups were observed.

#### Results from regions-of-Interest (ROI) analysis

We focused our ROI analyses on the predefined ROIs, including the amygdala, VS, putamen, and vmPFC, and examined whether neural responses during the contrasts of interest were associated with AUD status, AUDIT scores, and the corresponding behavioural PIT measures. For the alcohol-specific PIT cue contrast (alcohol CS vs. juice CS), the AUD group showed a trend-level stronger differential response to alcohol relative to juice cues in the amygdala compared with the non-AUD group (β = 0.218, SE = 0.115, t(160) = 1.91, p = .059, η² = .022, see Fig. 3C) and a trend-level association with AUDIT total score (β = 0.016, SE = 0.009, t(149) = 1.83, p = .069, η² = .022, see Fig. 3D). The association with AUDIT-C was significant, β = 0.056, SE = 0.022, t(149) = 2.51, p = .013, η² = .040. Given the left-lateralized amygdala responses observed at the whole-brain level (Figure 3B), we further examined the left amygdala, where responses were stronger in AUD, β = 0.267, SE = 0.131, t(160) = 2.04, p = .043, η² = .025 (Figure 3C), and positively associated with both AUDIT total, β = 0.025, SE = 0.010, t(149) = 2.48, p = .014, η² = .040 (Figure 3D), and AUDIT-C, β = 0.081, SE = 0.025, t(149) = 3.19, p = .002, η² = .064. No significant group differences were found for the other ROIs (all p > .210). The differential neural responses in the vmPFC showed a positive association with AUDIT-C (β = 0.038, SE = 0.016, t(149) = 2.35, p = .020, η² = .036), but not AUDIT total score (β = 0.009, SE = 0.006, t(149) = 1.39, p = .168, η² = .013). Neither putamen nor VS responses were not associated with AUDIT-C or AUDIT total score (all p > .119).

For the general PIT cue contrast, ROI analyses showed no significant group differences between AUD and non-AUD participants (all p > .103), and no significant associations with AUDIT total or AUDIT-C (all p > .159).

To test whether individual differences in behavioural PIT effects were reflected in the corresponding ROI responses, we examined the associations between behavioural specific and general PIT scores and neural responses in the selected ROIs using robust regressions. For specific PIT, we did not find any association between the behavioural score and the neural responses in any of the ROIs (all p > .113). For general PIT, robust regression showed significant positive associations between the behavioral general PIT score and the corresponding ROI responses in the amygdala, β = 0.100, SE = 0.059, p = .05, 95% CI [0.01, 0.29], putamen, β = 0.201, SE = 0.051, p = .001, CI [0.08, 0.27], and VS, β = 0.073, SE = 0.038, p = .006, CI [0.02, 0.17]. The association in the vmPFC was marginally positive, β = 0.065, SE = 0.041, p = .056, CI [−0.0002, 0.15].

A further exploratory ROI analysis of the anterior insula (aINS) was performed, motivated by its prominent engagement in the alcohol-versus-juice cue contrast and its proposed role in interoceptive/salience processes during alcohol cue exposure (49, 50). The aINS cluster was defined using cytoarchitectonic probabilistic maps from the JuBrain atlas (SPM Anatomy Toolbox; refs. 51, 52, 53), comprising subregions LD4, LD6, and LD7. The resulting mask was also resliced to the functional voxel size and binarized for ROI analyses. No group differences were found within the aINS. Nevertheless, aINS cue responses were positively associated with behavioral PIT parameters: alcohol-versus-juice responses were associated with specific PIT, β = 0.003, SE = 0.001, p = .007, 95% CI [0.001, 0.005], and responses to +€10 versus −€10 cues were associated with general PIT, β = 0.136, SE = 0.047, p = .002, CI [0.056, 0.232].

## Discussion

Using a novel full-transfer PIT task with gustatory alcohol rewards, we found that the alcohol-specific PIT effect was stronger in participants with AUD than in participants without AUD. It is worth noting that participants in the AUD sample had mostly mild-to-moderate AUD and were not treatment seeking. The alcohol-specific PIT effect was further associated with dimensional measures of AUD severity (number of AUD criteria and AUDIT score), alcohol use and craving. Importantly, it was also consistently associated with AUD severity and alcohol consumptions during the one-year follow-up. Additionally, AUD-related differences were also observed at the neural level: participants with AUD also showed enhanced fMRI responses in the left amygdala during alcohol-cue relative to juice-cue presentation, and these responses were also positively associated with AUDIT scores. In contrast, the general PIT effect elicited by monetary rewards was negatively associated with AUDIT total score but not with AUD diagnosis or other dimensional measures.

To our knowledge, this is the first study to provide empirical evidence in humans of an association between alcohol-specific PIT effects and AUD diagnosis in a clinical sample. Given its sensitivity to alcohol-related measures, alcohol-specific PIT may provide a candidate mechanistic marker for identifying at-risk individuals and evaluating the effects of interventions, complementing subjective measures such as craving (54) with a quantifiable index of cue-driven alcohol-seeking behavior.

The association between alcohol-specific PIT and AUD is consistent with PIT accounts proposing that reward-predictive cues can bias instrumental action selection, including cue-congruent choice tendencies in outcome-specific PIT (e.g., 2), and aligns with cue-reactivity evidence that alcohol-related cues can evoke motivational responses linked to alcohol use (55, 56). Our task elicited a strong alcohol-specific PIT effect overall (effect size: r = 0.70), indicating that alcohol and juice cues presented in our task systematically guided instrumental choice across participants. The stronger alcohol-specific PIT effect in the AUD group may indicate that alcohol cues more strongly guide outcome-specific choice in AUD, resulting in a more pronounced shift toward alcohol-associated actions, particularly in the early stages of addiction (Hogarth et al., 2012).In everyday life, alcohol cues are common and difficult to avoid. Repeated cue-driven shifts toward alcohol-associated choice may contribute to higher alcohol exposure in contexts where alcohol cues are frequently encountered, particularly when combined with alcohol’s reinforcing effects (57, 58). Consistent with this interpretation, specific PIT was associated not only with baseline drinking behavior but also with higher AUD severity and consumption across the one-year follow-up.

Contrary to our initial hypothesis that the general PIT effect would be increased in AUD, general PIT was not associated with AUD group and even showed a negative association with AUDIT total scores, as well as a similar but weaker association with AUDIT-C. As our full-transfer task design enabled us to evaluate both alcohol-specific and general PIT within the same paradigm, these divergent associations with AUD suggest that the two types of transfer may reflect different ways in which Pavlovian cues influence instrumental behavior. This is consistent with the view that these two forms of transfer involve different neural circuits. Animal lesion studies suggest that specific and general PIT rely on partially distinct amygdala–striatal circuits (2). Overall, while the present findings support a distinction between alcohol-specific and general PIT, the unexpected negative AUDIT association for general PIT should be interpreted with caution. Future research is required to establish whether this pattern can be replicated and whether it depends on the reward modality used to elicit general PIT.

Our imaging findings may help explain why alcohol use tends to be repeated. Following gustatory conditioning, alcohol cues elicited stronger responses in the left insula, bilateral precuneus, and temporo-parietal association cortex, consistent with engagement of salience/interoceptive processing (49, 50), together with broader multimodal associative processing, and memory-related processes (59–63). Compared with previous studies that relied primarily on visual alcohol or food pictures (e.g., 20), gustatory conditioning may recruit a broader and potentially more behaviorally relevant representation of alcohol-related cues, contributing to the strong specific PIT effect observed in the present study. More broadly, this richer cue representation may help explain why alcohol cues can be particularly effective in promoting alcohol-seeking behavior in everyday settings.

In contrast, juice cues elicited stronger left amygdala responses than alcohol cues. Given that participants generally preferred juice and chose it more often across conditions (67% juice choices), this pattern may reflect the amygdala’s sensitivity to the motivational relevance of outcomes in a context-dependent manner (64, 65). This finding also fits with previous imaging studies implicating the role of the amygdala in specific PIT (27, 28, 30). Importantly, the differential left amygdala responses to alcohol cues relative to juice cues were greater in AUD and were positively associated with AUDIT total and AUDIT-C scores. This suggests that, with increasing alcohol-related problems, the amygdala may encode greater motivational value of alcohol cues. The left-lateralized amygdala responses are broadly consistent with accounts linking the left amygdala to more sustained processing and the right amygdala to faster, more automatic processing (Ocklenburg et al., 2022; Sergerie et al., 2008). However, these responses were not directly associated with behavioral alcohol-specific PIT, making the interpretation less straightforward. This suggests that stronger amygdala responses to alcohol cues in AUD may not directly explain individual differences in cue-guided alcohol choice. In contrast, between-subject alcohol-specific PIT was associated with ACC and postcentral activity, consistent with increased monitoring and sensorimotor engagement during cue-guided action selection (66).

In contrast to the specific PIT effect, the cue effect for general PIT was associated with a broader neural network, including the vmPFC, putamen, posterior cingulate cortex (PCC), and thalamus. This pattern may reflect more distributed value-related processing and is broadly consistent with the involvement of these regions in valuation and reward circuitry (67, 68). Individual differences in association with the behavioural general PIT effect yielded an even broader neural pattern, extending to dmPFC, anterior cingulate cortex (ACC) as well as insula. The broader pattern indicates that individuals showing stronger general PIT may recruit not only valuation circuitry but also monitoring/control and salience processes during cue-guided action selection (50, 69–72). Compared with the more modest behavioral effects, which may be limited by the response-rate index (e.g., an upper bound on how many button presses can be made within limited time), neural responses in valuation and monitoring/salience systems may provide a more sensitive measure of cue susceptibility and cue-guided action selection.

Smoking status was not associated with either alcohol-specific or general PIT. This null finding for alcohol-specific PIT is not unexpected, since the effect was defined by outcome-specific choice between alcohol– and juice-related cues, and therefore did not directly involve smoking-related cues or outcomes. Previous studies on specific PIT using smoking-related cues in smokers have also reported mixed results (26, 73, 74), see also Garbusow et al. (75) for a review. This may be due in part to the limited potency of cues in these paradigms, which rely mainly on pictorial smoking cues. Future studies may test this more directly by using more ecologically valid smoking cues. For general PIT, however, we had expected smoking status to be associated with a stronger reward-related response vigor, but this was not supported, except for a weak interaction with AUD status. Given the limited evidence on the association between general PIT and smoking, this possible relationship requires future investigation. Finally, our findings here add to the accumulating evidence linking PIT effects to AUD and high-risk drinking in single-lever PIT paradigms (14, 16, 17, 19). Although further work is needed to clarify the underlying mechanisms, the converging evidence supports PIT-related cue control as a meaningful process in alcohol addiction.

## Limitations

The AUD and non-AUD groups also differed in age, which was not intended during recruitment. However, age was not significantly associated with the main outcomes, and age-adjusted models yielded consistent results. Another methodological consideration concerns the different reward modalities used for the two PIT measures: general PIT was indexed using monetary cues, whereas alcohol-specific PIT relied on gustatory alcohol rewards. As discussed in Belanger et al. (24), differences in reward modality and subjective value may influence PIT magnitude and limit direct comparability between these effects. The broad definition of occasional smoking represents a further limitation, as it may have combined participants with markedly different smoking patterns, ranging from social smoking to regular non-daily smoking and recent smoking cessation. This heterogeneity may have reduced sensitivity to detect smoking-related effects on general PIT, whereas no specific association between smoking status and alcohol-specific PIT was hypothesized a priori. In addition, follow-up participation was limited, reducing statistical power for longitudinal analyses and potentially introducing attrition bias if dropout was non-random. Finally, the neural results reported at the whole-brain level were uncorrected and should therefore be interpreted with caution. In addition, the multiband fMRI acquisition and associated reductions in subcortical temporal signal-to-noise (76) may have reduced sensitivity in ventral striatal regions, which could have limited our ability to detect ventral striatal associations.

## Conclusions

Using a PIT paradigm with trial-by-trial gustatory reward delivery during learning, we found a stronger alcohol-specific PIT effect in participants with AUD. Across the full sample, greater alcohol-specific PIT was also cross-sectionally associated with higher AUD severity and alcohol consumption at baseline, and these associations extended across the one-year follow-up period. This behavioral pattern was accompanied by greater differential left amygdala responses to alcohol relative to juice cues in participants with AUD. In contrast, general PIT robustly engaged reward-related neural systems but was unrelated to AUD status; instead, it was negatively associated with AUDIT total scores. These findings suggest that in mild-to-moderate, non-treatment-seeking AUD, the critical motivational bias may not reflect a generalized enhancement of Pavlovian influence, but a selective alcohol-specific bias in cue-elicited action selection. As a quantifiable behavioural index of cue-driven alcohol seeking, alcohol-specific PIT may therefore provide a sensitive mechanistic marker for identifying at-risk individuals and evaluating interventions aimed at reducing alcohol cue-triggered choice biases before they contribute to more persistent alcohol-seeking behaviour.

## Acknowledgments and Disclosures

This study was supported by the German Research Foundation (Deutsche Forschungsgemeinschaft) (Grant Nos. 402170461 [TRR 265], 454245598 [IRTG 2773], and 460922097 [INST 269/881-1 FUGG]). We thank Angela Hentschel for her support in participant recruitment and data collection, and Dr. Fabian Arntz for his important contribution to the organization of the longitudinal drinking data. ChatGPT was used to generate the pump, speaker and hand icons in Figure 1 and 2. The authors reviewed the AI-generated content and take full responsibility for it.

## Notes

### Competing Interest Statement

The authors have declared no competing interest.

## References

1. Holmes NM, Marchand AR, Coutureau E (2010): Pavlovian to instrumental transfer: a neurobehavioural perspective. Neurosci Biobehav Rev. 34:1277–1295.

2. Cartoni E, Balleine B, Baldassarre G (2016): Appetitive Pavlovian-instrumental Transfer: A review. Neurosci Biobehav Rev. 71:829–848.

3. Corbit LH, Balleine BW (2011): The general and outcome-specific forms of Pavlovian-instrumental transfer are differentially mediated by the nucleus accumbens core and shell. J Neurosci. 31:11786–11794.

4. Corbit LH, Balleine BW (2005): Double dissociation of basolateral and central amygdala lesions on the general and outcome-specific forms of pavlovian-instrumental transfer. J Neurosci. 25:962–970.

5. Mahlberg J, Seabrooke T, Weidemann G, Hogarth L, Mitchell CJ, Moustafa AA (2021): Human appetitive Pavlovian-to-instrumental transfer: a goal-directed account. Psychol Res. 85:449–463.

6. Seabrooke T, Hogarth L, Edmunds CER, Mitchell CJ (2019): Goal-directed control in Pavlovian-instrumental transfer. J Exp Psychol Anim Learn Cogn. 45:95–101.

7. Seabrooke T, Hogarth L, Mitchell CJ (2016): The propositional basis of cue-controlled reward seeking. Q J Exp Psychol (Hove). 69:2452–2470.

8. Seabrooke T, Le Pelley ME, Hogarth L, Mitchell CJ (2017): Evidence of a goal-directed process in human Pavlovian-instrumental transfer. J Exp Psychol Anim Learn Cogn. 43:377–387.

9. Seabrooke T, Le Pelley ME, Porter A, Mitchell CJ (2018): Extinguishing cue-controlled reward choice: Effects of Pavlovian extinction on outcome-selective Pavlovian-instrumental transfer. J Exp Psychol Anim Learn Cogn. 44:280–292.

10. Berridge KC, Robinson TE (2016): Liking, wanting, and the incentive-sensitization theory of addiction. Am Psychol. 71:670–679.

11. Robinson TE, Berridge KC (1993): The neural basis of drug craving: an incentive-sensitization theory of addiction. Brain Res Brain Res Rev. 18:247–291.

12. Robinson TE, Berridge KC (2008): The incentive sensitization theory of addiction: some current issues. Philos T R Soc B. 363:3137–3146.

13. Hogarth L (2012): Goal-directed and transfer-cue-elicited drug-seeking are dissociated by pharmacotherapy: evidence for independent additive controllers. J Exp Psychol Anim Behav Process. 38:266–278.

14. Chen H, Nebe S, Mojtahedzadeh N, Kuitunen-Paul S, Garbusow M, Schad DJ, et al. (2021): Susceptibility to interference between Pavlovian and instrumental control is associated with early hazardous alcohol use. Addiction biology.e12983.

15. Garbusow M, Nebe S, Sommer C, Kuitunen-Paul S, Sebold M, Schad DJ, et al. (2019): Pavlovian-To-Instrumental Transfer and Alcohol Consumption in Young Male Social Drinkers: Behavioral, Neural and Polygenic Correlates. J Clin Med. 8:1188.

16. Garbusow M, Schad DJ, Sebold M, Friedel E, Bernhardt N, Koch SP, et al. (2016): Pavlovian-to-instrumental transfer effects in the nucleus accumbens relate to relapse in alcohol dependence. Addict Biol. 21:719–731.

17. Sommer C, Birkenstock J, Garbusow M, Obst E, Schad DJ, Bernhardt N, et al. (2020): Dysfunctional approach behavior triggered by alcohol-unrelated Pavlovian cues predicts long-term relapse in alcohol dependence. Addict Biol. 25:e12703.

18. Sommer C, Garbusow M, Junger E, Pooseh S, Bernhardt N, Birkenstock J, et al. (2017): Strong seduction: impulsivity and the impact of contextual cues on instrumental behavior in alcohol dependence. Transl Psychiatry. 7:e1183.

19. Chen K, Schlagenhauf F, Sebold M, Kuitunen-Paul S, Chen H, Huys QJ, et al. (2023): The association of non–drug-related Pavlovian-to-instrumental transfer effect in nucleus accumbens with relapse in alcohol dependence: A replication. Biological Psychiatry. 93:558–565.

20. Martinovic J, Jones A, Christiansen P, Rose AK, Hogarth L, Field M (2014): Electrophysiological responses to alcohol cues are not associated with Pavlovian-to-instrumental transfer in social drinkers. PloS one. 9:e94605.

21. Hardy L, Mitchell C, Seabrooke T, Hogarth L (2017): Drug cue reactivity involves hierarchical instrumental learning: evidence from a biconditional Pavlovian to instrumental transfer task. Psychopharmacology. 234:1977–1984.

22. Mahlberg J, Weidemann G, Hogarth L, Moustafa AA (2019): Cue-elicited craving and human Pavlovian-to-instrumental transfer. Addiction Research & Theory. 27:482–488.

23. van Timmeren T, Quail SL, Balleine BW, Geurts DE, Goudriaan AE, van Holst RJ (2020): Intact corticostriatal control of goal-directed action in Alcohol Use Disorder: a Pavlovian-to-instrumental transfer and outcome-devaluation study. Sci Rep. 10:1–12.

24. Belanger MJ, Chen H, Hentschel A, Garbusow M, Ebrahimi C, Knorr FG, et al. (2022): Development of novel tasks to assess outcome-specific and general Pavlovian-to-instrumental transfer in humans. Neuropsychobiology. 81:370–386.

25. Manglani HR, Lewis AH, Wilson SJ, Delgado MR (2017): Pavlovian-to-instrumental transfer of nicotine and food cues in deprived cigarette smokers. Nicotine & Tobacco Research. 19:670–676.

26. Hogarth L, Chase HW (2012): Evaluating psychological markers for human nicotine dependence: tobacco choice, extinction, and Pavlovian-to-instrumental transfer. Experimental and clinical psychopharmacology. 20:213.

27. Prevost C, Liljeholm M, Tyszka JM, O’Doherty JP (2012): Neural correlates of specific and general Pavlovian-to-Instrumental Transfer within human amygdalar subregions: a high-resolution fMRI study. J Neurosci. 32:8383–8390.

28. van Steenbergen H, Watson P, Wiers RW, Hommel B, de Wit S (2017): Dissociable corticostriatal circuits underlie goal-directed vs. cue-elicited habitual food seeking after satiation: evidence from a multimodal MRI study. Eur J Neurosci. 46:1815–1827.

29. Lewis AH, Niznikiewicz MA, Delamater AR, Delgado MR (2013): Avoidance-based human Pavlovian-to-instrumental transfer. Eur J Neurosci. 38:3740–3748.

30. Morris RW, Quail S, Griffiths KR, Green MJ, Balleine BW (2015): Corticostriatal control of goal-directed action is impaired in schizophrenia. Biological psychiatry. 77:187–195.

31. Heinz A, Kiefer F, Smolka MN, Endrass T, Beste C, Beck A, et al. (2020): Addiction Research Consortium: Losing and regaining control over drug intake (ReCoDe)—From trajectories to mechanisms and interventions. Addiction Biology. 25:e12866.

32. American Psychiatric Association (2013): Diagnostic and statistical manual of mental disorders. 5th ed. Arlington, VA: American Psychiatric Publishing.

33. Beesdo-Baum K, Zaudig M, Wittchen H (2019): SCID-5-CV Strukturiertes Klinisches Interview für DSM-5®–Störungen. Göttingen: Hogrefe(Deutsche Bearbeitung des Structured Clinical Interview for DSM-5®–Clinician Version von Michael B First, Janet BW Williams, Rhonda S Karg, Robert L Spitzer).

34. Kuitunen-Paul S, Rehm, J., Lachenmeier, D. W., Kadrić, F., Kuitunen, P. T., Wittchen, H. U., & Manthey, J. (2017): Assessment of alcoholic standard drinks using the Munich composite international diagnostic interview (M – CIDI): An evaluation and subsequent revision. International journal of methods in psychiatric research. 26:e1563.

35. Saunders JB, Aasland OG, Babor TF, De la Fuente JR, Grant M (1993): Development of the alcohol use disorders identification test (AUDIT): WHO collaborative project on early detection of persons with harmful alcohol consumption – II. Addiction. 88:791–804.

36. Vollstädt-Klein S, Leménager T, Jorde A, Kiefer F, Nakovics H (2015): Development and validation of the craving automated scale for alcohol. Alcohol Clin Exp Res. 39:333–342.

37. Heatherton TF, Kozlowski LT, Frecker RC, Fagerstrom KO (1991): The Fagerstrom Test for Nicotine Dependence – a Revision of the Fagerstrom Tolerance Questionnaire. Brit J Addict. 86:1119–1127.

38. Höfler M (2026): Robust tests should be the default, not the backup. Peer Community Journal. 6.

39. Venables W, Ripley B (2002): Modern Applied Statistics with S.

40. Koller M (2016): robustlmm: an R package for robust estimation of linear mixed-effects models. Journal of statistical software. 75:1–24.

41. Lund TE, Nørgaard MD, Rostrup E, Rowe JB, Paulson OB (2005): Motion or activity: their role in intra-and inter-subject variation in fMRI. Neuroimage. 26:960–964.

42. Power JD, Barnes KA, Snyder AZ, Schlaggar BL, Petersen SE (2012): Spurious but systematic correlations in functional connectivity MRI networks arise from subject motion. Neuroimage. 59:2142–2154.

43. Siegel JS, Power JD, Dubis JW, Vogel AC, Church JA, Schlaggar BL, et al. (2014): Statistical improvements in functional magnetic resonance imaging analyses produced by censoring high – motion data points. Hum Brain Mapp. 35:1981–1996.

44. Friston KJ, Williams S, Howard R, Frackowiak RS, Turner R (1996): Movement – related effects in fMRI time – series. Magnetic resonance in medicine. 35:346–355.

45. Tzourio-Mazoyer N, Landeau B, Papathanassiou D, Crivello F, Etard O, Delcroix N, et al. (2002): Automated anatomical labeling of activations in SPM using a macroscopic anatomical parcellation of the MNI MRI single-subject brain. Neuroimage. 15:273–289.

46. Maldjian JA, Laurienti PJ, Burdette JH (2004): Precentral gyrus discrepancy in electronic versions of the Talairach atlas. Neuroimage. 21:450–455.

47. Maldjian JA, Laurienti PJ, Kraft RA, Burdette JH (2003): An automated method for neuroanatomic and cytoarchitectonic atlas-based interrogation of fMRI data sets. Neuroimage. 19:1233–1239.

48. Nielsen FA, Hansen LK (2002): Automatic anatomical labeling of Talairach coordinates and generation of volumes of interest via the BrainMap database. Neuroimage. 16:1126–1128.

49. Simmons WK, Avery JA, Barcalow JC, Bodurka J, Drevets WC, Bellgowan P (2013): Keeping the body in mind: insula functional organization and functional connectivity integrate interoceptive, exteroceptive, and emotional awareness. Hum Brain Mapp. 34:2944–2958.

50. Uddin LQ (2015): Salience processing and insular cortical function and dysfunction. Nat Rev Neurosci. 16:55–61.

51. Eickhoff SB, Heim S, Zilles K, Amunts K (2006): Testing anatomically specified hypotheses in functional imaging using cytoarchitectonic maps. Neuroimage. 32:570–582.

52. Eickhoff SB, Paus T, Caspers S, Grosbras M-H, Evans AC, Zilles K, et al. (2007): Assignment of functional activations to probabilistic cytoarchitectonic areas revisited. Neuroimage. 36:511–521.

53. Eickhoff SB, Stephan KE, Mohlberg H, Grefkes C, Fink GR, Amunts K, et al. (2005): A new SPM toolbox for combining probabilistic cytoarchitectonic maps and functional imaging data. Neuroimage. 25:1325–1335.

54. Kavanagh DJ, Statham DJ, Feeney GF, Young RM, May J, Andrade J, et al. (2013): Measurement of alcohol craving. Addictive behaviors. 38:1572–1584.

55. Schacht JP, Anton RF, Myrick H (2013): Functional neuroimaging studies of alcohol cue reactivity: a quantitative meta-analysis and systematic review. Addict Biol. 18:121–133.

56. Zeng J, Yu S, Cao H, Su Y, Dong Z, Yang X (2021): Neurobiological correlates of cue-reactivity in alcohol-use disorders: A voxel-wise meta-analysis of fMRI studies. Neurosci Biobehav Rev. 128:294–310.

57. Koob GF, Volkow ND (2016): Neurobiology of addiction: a neurocircuitry analysis. Lancet Psychiatry. 3:760–773.

58. Heinz A, Beck A, Halil MG, Pilhatsch M, Smolka MN, Liu S (2019): Addiction as learned behavior patterns. J Clin Med. 8:1086.

59. Bonnici HM, Richter FR, Yazar Y, Simons JS (2016): Multimodal feature integration in the angular gyrus during episodic and semantic retrieval. J Neurosci. 36:5462–5471.

60. Tibon R, Fuhrmann D, Levy DA, Simons JS, Henson RN (2019): Multimodal integration and vividness in the angular gyrus during episodic encoding and retrieval. J Neurosci. 39:4365–4374.

61. Dahl CD, Logothetis NK, Kayser C (2009): Spatial organization of multisensory responses in temporal association cortex. J Neurosci. 29:11924–11932.

62. James TW, VanDerKlok RM, Stevenson RA, James KH (2011): Multisensory perception of action in posterior temporal and parietal cortices. Neuropsychologia. 49:108–114.

63. Dadario NB, Sughrue ME (2023): The functional role of the precuneus. Brain. 146:3598–3607.

64. Cunningham WA, Van Bavel JJ, Johnsen IR (2008): Affective flexibility: evaluative processing goals shape amygdala activity. Psychol Sci. 19:152–160.

65. Janak PH, Tye KM (2015): From circuits to behaviour in the amygdala. Nature. 517:284–292.

66. Fan J, Hof PR, Guise KG, Fossella JA, Posner MI (2008): The functional integration of the anterior cingulate cortex during conflict processing. Cereb Cortex. 18:796–805.

67. Bartra O, McGuire JT, Kable JW (2013): The valuation system: a coordinate-based meta-analysis of BOLD fMRI experiments examining neural correlates of subjective value. Neuroimage. 76:412–427.

68. Haber SN, Knutson B (2010): The reward circuit: linking primate anatomy and human imaging. Neuropsychopharmacology. 35:4–26.

69. Taren AA, Venkatraman V, Huettel SA (2011): A parallel functional topography between medial and lateral prefrontal cortex: evidence and implications for cognitive control. J Neurosci. 31:5026–5031.

70. de Kloet SF, Bruinsma B, Terra H, Heistek TS, Passchier EM, van den Berg AR, et al. (2021): Bi-directional regulation of cognitive control by distinct prefrontal cortical output neurons to thalamus and striatum. Nat Commun. 12:1994.

71. Botvinick MM, Cohen JD, Carter CS (2004): Conflict monitoring and anterior cingulate cortex: an update. Trends Cogn Sci. 8:539–546.

72. Billeke P, Ossandon T, Perrone-Bertolotti M, Kahane P, Bastin J, Jerbi K, et al. (2020): Human anterior insula encodes performance feedback and relays prediction error to the medial prefrontal cortex. Cereb Cortex. 30:4011–4025.

73. Hogarth L, Dickinson A, Wright A, Kouvaraki M, Duka T (2007): The role of drug expectancy in the control of human drug seeking. J Exp Psychol Anim Behav Process. 33:484–496.

74. Hogarth L, Field M, Rose AK (2013): Phasic transition from goal-directed to habitual control over drug-seeking produced by conflicting reinforcer expectancy. Addict Biol. 18:88–97.

75. Garbusow M, Ebrahimi C, Riemerschmid C, Daldrup L, Rothkirch M, Chen K, et al. (2022): Pavlovian-to-Instrumental Transfer across Mental Disorders: A Review. Neuropsychobiology. 81:418–437.

76. Srirangarajan T, Mortazavi L, Bortolini T, Moll J, Knutson B (2021): Multi-band FMRI compromises detection of mesolimbic reward responses. Neuroimage. 244:118617.

