## Supplementary Material for "Enhanced Alcohol-Specific but not General Pavlovian-to-Instrumental Transfer in Alcohol Use Disorder"

### S-1: Smoking Quantity-Frequency Questionnaire

Tobacco and e-cigarette use were assessed using an interviewer-administered quantity/frequency questionnaire. The original German questionnaire used “tobacco” as a collective term referring to all tobacco products used by the participant.

**Item 1: Type of tobacco or nicotine product used**

Participants were asked: “What did you smoke/use?” Multiple responses were allowed. Response options were: cigarettes, cigarillos, cigars, water pipe/hookah (shisha), e-cigarettes/liquids, heated tobacco products, snuff tobacco, and other tobacco products.

**Item 2: Smoking/use frequency during the past 3 months**

Participants were asked: “On how many days during the past 3 months did you smoke at least one cigarette, or use an e-cigarette or tobacco product at least once?” Responses were entered as a number between 0 and 93.

**Item 3: Time scale of frequency report**

Participants were asked: “What format was used for reporting the number of days in the previous question?” Response options were: number of days during the past 3 months, or number of days per week within the past 3 months.

**Item 4: Maximum recent smoking/use quantity**

Participants were asked: “During the period in the past 3 months when you smoked/used the most, how many cigarettes did you smoke on average per day, or how often did you use an e-cigarette on average per day?” Responses were entered as a number between 1 and 999.

**Item 5: Previous smoking cessation programme**

Participants were asked: “Have you ever participated in a smoking cessation programme, for example using nicotine patches, behavioural therapy, or a similar intervention?” Response options were yes or no.

**Item 6: Lifetime regular smoking/use**

Participants were asked: “Have you ever smoked regularly in your lifetime, defined as smoking at least one cigarette, or using another tobacco product, daily for a period of 4 weeks or longer?” Response options were yes or no.

**Item 7: Age at first regular smoking/use**

If lifetime regular smoking/use was endorsed, participants were asked: “At what age did you first smoke at least one cigarette, or use an e-cigarette, daily for a period of more than 4 weeks?” The interviewer entered the participant’s age in years.

**Item 8: Age at last regular smoking/use**

If lifetime regular smoking/use was endorsed, participants were asked: “At what age did you last smoke at least one cigarette, or use an e-cigarette, daily for a period of more than 4 weeks?” The interviewer entered the participant’s age in years.

**Item 9: Maximum lifetime smoking/use quantity**

If lifetime regular smoking/use was endorsed, participants were asked: “Please think of the period in your life during which you smoked/used the most. How many cigarettes did you smoke on average per day, or how often did you use an e-cigarette on average per day?” Responses were entered as a number between 1 and 999.

### S-2: Analysis of query trials

To assess explicit cue–outcome knowledge, participants completed a query phase after the transfer phase. In this phase, each of the five Pavlovian cues was presented five times. On each trial, participants selected the outcome associated with the cue from multiple response options. Query accuracy was calculated as the proportion of correctly identified cue–outcome associations. We examined whether individual differences in query accuracy were associated with behavioural specific or general PIT effects.

90% of all participants have an accuracy of 100% during the query trials. There was no association between either specific or general PIT effect and the accuracy during the query trials (p > 0.764; Figure S1). There were three participants who had an accuracy below 50%, and the effect size for both specific and general PIT improved slightly after excluding the three participants (specific PIT: from 0.702 to 0.714; general PIT: 0.357 to 0.396). Given the generally high query performance across participants, and because explicit cue–outcome knowledge and cue-driven behavioural influence during PIT may reflect partly distinct processes, we did not conduct further analyses based on query accuracy.

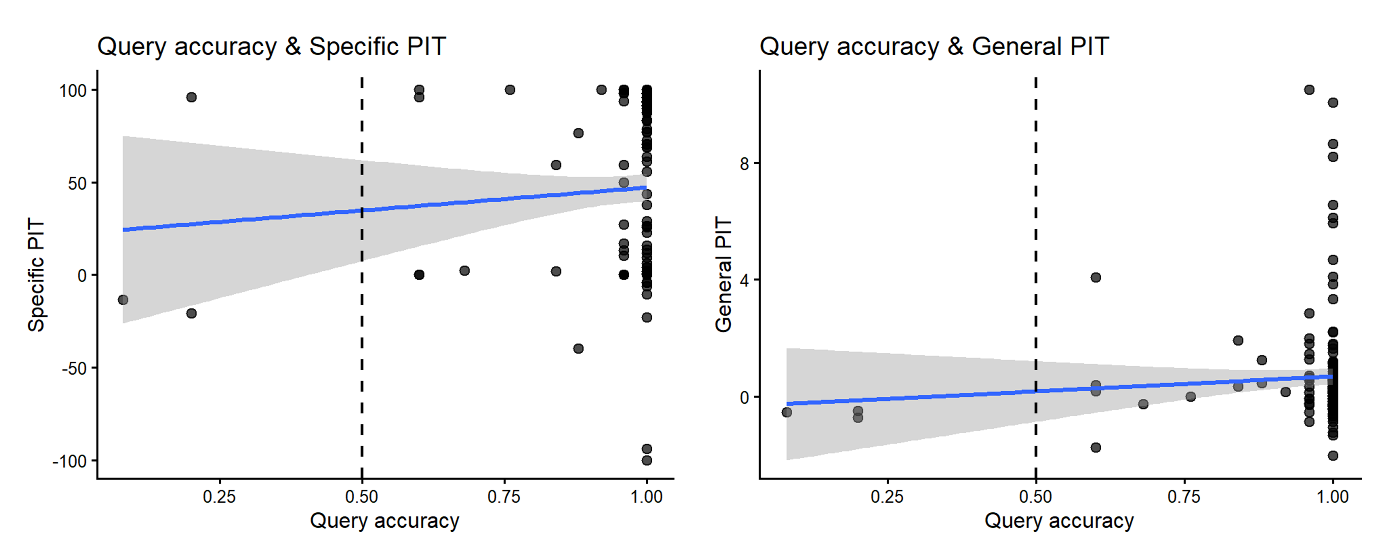

**Figure S1: Association between query accuracy and specific PIT scores (left) and general PIT scores (right).** Query accuracy was not associated with either specific or general PIT. The dashed vertical line indicates 50% query accuracy. Blue lines indicate linear regression fits with 95% confidence intervals.

### S-3: Behaviour (choice or vigor)-modulated neural PIT contrasts

In the main analyses, we focused on between-subject associations between behavioural PIT scores and neural responses to the corresponding Pavlovian cues. As an additional analysis, we examined whether trial-wise choice and vigor-modulated PIT contrasts were associated with neural responses. These analyses were included to assess whether cue-dependent behavioural variation within individuals was reflected in brain activation. However, they should be interpreted as exploratory and complementary to the main analyses, because choice-/vigor-modulated regressors depend on sufficient trial-wise behavioural variability and may therefore be less stable than between-subject PIT scores.

For the choice-modulated specific PIT effect (alcohol vs. juice choice; contrast of the parametric regressors), no cerebral gray-matter (cortical or subcortical) responses survived this threshold; only two clusters were observed in bilateral cerebellum (see Table S1). For the opposite parametric contrast (i.e., greater responses for juice-over-alcohol choice), neural responses were observed in a left-lateralized fronto-parietal sensorimotor network, including the anterior supramarginal gyrus, precentral gyrus, and superior frontal gyrus (see Table S1). However, this analysis was based on only 68 participants with complete data, as not all participants made both alcohol and juice choices during alcohol and juice CS presentations.

In contrast, for vigor-modulated general PIT effect, using the same whole-brain threshold, the within-subject general PIT effect also showed no significant clusters.

Overall, these supplementary findings suggest that choice-/vigor-modulated neural PIT effects were weaker and less robust than cue-evoked responses and between-subject PIT associations. For the within-subject alcohol-specific PIT modulator (alcohol choice higher under alcohol vs. juice cues), the cue-congruent contrast was confined to bilateral cerebellar Crus, consistent with cerebellar contributions to reinforcement learning and translating cue-related expectations into action selection (Kruithof et al., 2023). The reverse contrast (alcohol choice higher under juice vs. alcohol cues) recruited a left-lateralized fronto-parietal sensorimotor network, consistent with cue–action mapping and response preparation (Pellegrino et al., 2018), but not sufficiently process-specific to distinguish valuation from attentional or motoric contributions to the choice bias. This pattern is broadly consistent with prior work showing weaker within-subject than between-subject PIT effects (van Timmeren et al., 2020). Importantly, the alcohol-specific within-subject regressor was estimable only in participants with sufficient free-choice trial variability (n = 68), which limits power and generalizability. More generally, within-subject parametric effects may be attenuated by restricted trial-wise behavioural variance (e.g., bounded response rates or repeated identical choices) and by variance already captured by the cue-onset regressor.

### Table S1: Whole-brain analysis results

| **Contrast** | **Cluster** | **Cluster pFWE** | **Size** | **Region** | **Peak MNI** | | | **Peak-level t score** |
| --- | --- | --- | --- | --- | --- | --- | --- | --- |
|  |  |  |  |  | **x** | **y** | **z** |  |
| Juice > Alcohol | 1 | .363 | 80 | L Amygdala | -23 | -9 | -15 | 4.07 |
| Alcohol > Juice | 1 | <.001 | 2022 | R Superior Frontal Gyrus | 11 | 18 | 62 | 6.44 |
|  | 2 | <.001 | 869 | R Angular Gyrus (extending into posterior Supramarginal Gyrus) | 49 | -54 | 33 | 5.76 |
|  | 3 | <.001 | 815 | L Lateral Occipital Cortex | -59 | -64 | 36 | 5.71 |
|  | 4 | <.001 | 617 | R Precuneuous Cortex | 8 | -66 | 38 | 5.42 |
|  | 5 | <.001 | 554 | L Insular Cortex | -37 | 15 | 2 | 4.47 |
|  | 6 | <.001 | 552 | R Frontal Orbital Cortex (extending to Frontal Operculum Cortex) | 42 | 22 | -3 | 4.79 |
|  | 7 | .039 | 208 | R Frontal Pole | 18 | 49 | 33 | 4.24 |
|  | 8 | .066 | 176 | R Middle Frontal Gyrus | 40 | 20 | 45 | 4.22 |
|  | 9 | .072 | 171 | R Cingulate Gyrus, posterior division | 1 | -21 | 33 | 4.88 |
|  | 10 | .095 | 155 | L Middle Temporal Gyrus, posterior division | -56 | -33 | -8 | 4.32 |
|  | 11 | .350 | 82 | L Middle Frontal Gyrus | -37 | 15 | 43 | 4.12 |
|  | 12 | .369 | 79 | R Middle Temporal Gyrus, posterior division | 49 | -38 | -3 | 3.91 |
|  | 13 | .383 | 77 | R Middle Temporal Gyrus, posterior division | 56 | -11 | -27 | 4.18 |
| Alcohol > Juice choice parametric | 1 | .539 | 55 | R Cerebellum Crus | 37 | -78 | -24 | 3.51 |
|  | 2 | .579 | 51 | L Cerebellum Crus | -40 | -81 | -29 | 3.67 |
| Juice > Alcohol choice parametric | 1 | .021 | 227 | L Supramarginal Gyrus, anterior division | -61 | -26 | 31 | 3.96 |
|  | 2 | .100 | 141 | L Precentral Gyrus | -49 | 3 | 33 | 4.08 |
|  | 3 | .295 | 86 | L Superior Frontal Gyrus | -28 | -4 | 52 | 3.81 |
| Between-subject specific PIT effect (positive association) | 1 | .272 | 96 | L postcentral Gyrus | -61 | -16 | 31 | 4.19 |
|  | 2 | .426 | 71 | R Cingulate Gyrus, anterior division | 8 | 39 | 7 | 4.05 |
| Between-subject specific PIT effect (negative association) | 1 | .344 | 83 | R Lateral Occipital Cortex, inferior division | 49 | -78 | 7 | 3.86 |
| €+10 > €-10 cue contrast | 1 | <.001 | 17181 | L Putamen/striatum; posterior cingulate/precuneus; lateral occipital cortex; posterior temporal cortex (fusiform/ITG, planum temporale/STG); precentral/SMA; thalamus; angular gyrus; extends into medial temporal lobe (amygdala/hippocampus) | -30 | -6 | -10 | 5.99 |
|  | 2 | <.001 | 4063 | R Frontal Medial Cortex | 6 | 46 | -10 | 5.36 |
|  | 3 | .087 | 151 | L Frontal Orbitral Cortex | -47 | 27 | -17 | 4.6 |
|  | 4 | .111 | 138 | L Precentral Gyrus | -61 | 8 | 40 | 4.11 |
|  | 5 | .360 | 77 | R Cerebellum Crus | 44 | -76 | -36 | 4.41 |
|  | 6 | .412 | 70 | L Precentral Gyrus | -52 | -6 | 48 | 3.82 |
|  | 7 | .453 | 65 | R Temporal Pole | 42 | 15 | -29 | 4.17 |
|  | 8 | .535 | 56 | L Cerebellum VI | -23 | -64 | -22 | 3.79 |
| Between-subject general PIT effect (positive association) | 1 | <.001 | 43838 | L thalamus; cingulate cortex (anterior→posterior); basal ganglia (L putamen/pallidum; bilateral caudate; extends to R putamen; includes L amygdala); insula/operculum (parietal operculum); motor/premotor cortex (precentral; juxtapositional lobule/SMA); frontal cortex (L superior/middle frontal; R orbitofrontal; R frontal pole); brainstem | -8 | 3 | 38 | 9.27 |
| Between-subject general PIT effect (negative association) | 1 | .428 | 68 | R Cerebellum Crus | 13 | -90 | -22 | 4.97 |
|  | 2 | .564 | 53 | R Occipital Pole | 6 | -90 | 36 | 6.15 |

### Reference

Kruithof, E. S., Klaus, J., & Schutter, D. J. (2023). The human cerebellum in reward anticipation and outcome processing: An activation likelihood estimation meta-analysis. *Neuroscience & Biobehavioral Reviews*, *149*, 105171.

Pellegrino, G., Tomasevic, L., Herz, D. M., Larsen, K. M., & Siebner, H. R. (2018). Theta activity in the left dorsal premotor cortex during action re-evaluation and motor reprogramming. *Frontiers in Human Neuroscience*, *12*, 364.

van Timmeren, T., Quail, S. L., Balleine, B. W., Geurts, D. E., Goudriaan, A. E., & van Holst, R. J. (2020). Intact corticostriatal control of goal-directed action in Alcohol Use Disorder: a Pavlovian-to-instrumental transfer and outcome-devaluation study. *Scientific Reports*, *10*(1), 1-12. <https://doi.org/10.1038/s41598-020-61892-5>
